# Cortical Excitability during Fixations Drives Frequency-Specific Neural Activity in Children and Adults

**DOI:** 10.1101/2025.02.27.639836

**Authors:** I. Marriott Haresign, T. Charman, M.H. Johnson, L. Mason, T. Bazelmans, J. Begum-Ali, E.J.H. Jones, S.V. Wass

## Abstract

Extensive previous research has examined how neuronal oscillations support basic cognitive processes, from early development into adulthood. However, the question of how these oscillations originate and are maintained remains relatively underexplored. Here, we examine how transient increases in cortical excitability that occur time-locked to the offsets of spontaneous eye movements associate with frequency-specific neural activity, and how these relationships change over development and between contexts (social vs non-social). We examine two datasets of combined EEG and eye-tracking data from 24-month-old children (N=114) and adults (N=108) while they watched stimuli that were either social (an actor singing nursery rhythms) or non-social (dynamic toys). EEG data was time-locked to the offsets of eye movements and analysed using a spectrum of methods designed to highlight the progression of various neural signals across time, frequency, and space (topography). Fixation-related potentials (FRPs) manifest as a differentiable combination of eye movement-related artifact and genuine neural activity. Child FRPs are slower and unfold over longer time periods, which manifests as differences in the frequency domain. Even after removing artifact, dipoles associated with fixation-related P1 and N170 components manifest as Theta activity over fronto-central areas, along with activity in other frequencies, in children but not adults. Data sections where no fixation-related potentials are present show strongly attenuated oscillatory activity. Our results show that a variety of previously documented developmental effects in the frequency domain may be better understood as fine-grained, movement-induced brain states.

## 1. Introduction

During free behaviour, all animals sample from their environments periodically through behaviours including whisking, sniffing (Kleinfeld et al., 2016), and eye movements (Findlay & Gilchrist, 2003). These active exploratory behaviours occur most prominently in the 3-9Hz range (Haegens & Golumbic, 2018; Lakatos et al., 2019). In human adults, refoveating eye movements (known as saccades) take place at a modal interval of ∼300ms (3Hz) (Findlay & Walker, 1999; Leszczynski & Schroeder, 2019; Yarbus, 1967), but more fine-grained eye movements (known as micro-saccades) also occur within a fixation (Otero-Millan et al., 2008). Several studies have shown that eye movements are timed to consistent phases of neuronal oscillations, although effect sizes are small (Hogendoorn, 2016; Pan et al., 2021; Dimigen et al., 2011).

In humans, brief periods of cortical excitation follow fixations (known as fixation-related potentials (FRPs)), as the phases of neuronal oscillations transit from random to a highly organised state just after fixation onset (Maldonado et al., 2008). This transient phase of excitability following a fixation onset is known to be accompanied in adults by increased spectral power across lower frequency ranges (Dimigen et al., 2011; Leszczynski & Schroeder, 2019; Rajkai et al., 2008; Rousselet et al., 2007). There is some evidence in adults that FRPs differentiate between different viewing contexts. For example, when viewing direct vs averted eye gaze (Stephani et al., 2020) and incongruous objects in scenes (Coco et al., 2020). As yet, however, little work has investigated how these transient changes in cortical excitability contribute to the oscillatory activity documented both during unconstrained behaviour and experimental paradigms in the lab (Fries, 2023; Leszczynski & Schroeder, 2019; Rajkai et al., 2008; Yuval-Greenberg et al., 2008). In particular, no research has investigated how the relationship between transient changes in cortical excitability and oscillatory brain activity changes over development, and between different viewing contexts.

There is ample research to suggest that oscillatory activity observed during non-event-locked paradigms changes with age (Cellier et al., 2021; Stanyard et al., 2024; Wilkinson et al., 2024). For example, brain activity in younger participants is relatively more aperiodic (Stanyard et al., 2024; Wilkinson et al., 2024). Over time, changes occur in the relative proportions of frequency-specific ’periodic’ oscillations and the ’aperiodic signal’; and (relatedly) in 1/f power slope exponents (Voytek & Kramer, 2015; Marshall et al., 2002; Stanyard et al., 2024). Additionally, research has shown that the peak frequency (but not the power) of oscillatory brain activity tends to increase with age. Critically, infancy is characterized by a dominance of theta oscillations (3-6Hz) in posterior electrodes, whereas the peak frequency of dominant oscillations in the alpha range increases during early childhood (Cellier et al., 2021; Wilkinson et al., 2024). These changes are often considered to reflect the development of spontaneous (i.e., not task-evoked), intrinsic neural oscillations that are associated with changes in the structure and function of underlying brain networks (Cui et al., 2020; Marek et al., 2015; Wilkinson et al., 2024).

Paralleling these changes in oscillatory activity are changes in spontaneous eye movements. Ample evidence suggests that the micro-dynamics of eye movements change over development, showing both a decrease in modal fixation durations and a smaller proportion of long, extended fixations with age (Bronson, 1990, 1994; Luna et al., 2008). Developmental changes in eye movements also differ contingent on context. For example, when viewing dynamic vs static stimuli (Wass & Smith, 2014), or early vs late viewing of a static scene (Helo et al., 2016).

Research using task-evoked paradigms has also shown that the form and functionality of passive, event-related neural activity (ERPs) change over development (Oades et al., 1997). Passive, visual ERPs such as the P1 and N170 show reductions in latency and amplitude with age (Brecelj et al., 2002; Buchsbaum et al., 1974; Taylor et al., 2004). However, although parallel lines of research have shown that both ERP and frequency domain analyses show faster activity with age - i.e., decreases in latency and increases in the frequency of dominant oscillatory activity - no previous research has examined how changes in the time domain co-occur with neural activity in the frequency domain, i.e., whether it is better to think of these changes as two separate developmental mechanisms, or inter-related (although see Schneider & Maguire, 2018 for arguments in adults).

One framework which is relevant here is the debate over whether fixation- and event-related potentials result from evoked ‘additive’ brain activity (known as an evoked brain response) or induced changes in ongoing brain dynamics (known as an induced response) (Burgess, 2012; David et al., 2006; Makeig et al., 2002, 2004). Evoked responses are additive signals superimposed upon the background/ongoing EEG, whereas induced responses are changes in power and/or phase that take place within the background/ongoing EEG. Several researchers have, however, pointed out that these possibilities are non-exclusive (Burgess, 2012) and that methods for identifying phase changes around events struggle to disentangle the estimation of power and phase (Muthukumaraswamy et al., 2011; van Diepen & Mazaheri, 2018).

In addition to examining how the relationship between fixation-related potentials and underlying oscillatory brain activity changes over development, our secondary aim was to examine how this relationship changes between different viewing contexts. To examine this, we compared two viewing contexts, namely the presentation of video clips containing social vs non-social information.

We know that the human brain processes social and non-social stimuli differently. Research using task-evoked paradigms has shown greater N170 ERP amplitudes in response to faces vs objects and houses in adults (e.g., Bentin et al., 1996; Caldara et al., 2003; Carmel and Bentin, 2002; Itier et al., 2006; Itier and Taylor, 2004a,b; Rossion et al., 2000) and in infants (de Haan and Nelson, 1999; Guy et al., 2016; Peykarjou and Hoehl, 2013; Xie and Richards, 2016; Conte et al., 2020). These effects are, however, specific to the stage of the visual processing stream. Early stages of visual processing indexed by the P1 ERP components typically do not differentiate between social vs non-social viewing (Peykarjou and Hoehl, 2013), consistent with the consensus that the P1 reflects lower-level visual processing. Later-stage visual processing indexed by the Nc and P400 ERP components also do not reliably differentiate (see de Haan and Nelson, 1999 vs Guy et al., 2016), and seem more influenced by attention (Guy et al., 2016; Reynolds et al., 2010; Reynolds and Richards, 2005, 2009; Richards, 2003) and saliency (Courchesne et al., 1981; de Haan and Nelson, 1997, 1999; Guy et al., 2013; Reynolds et al., 2010; Richards, 2003; Webb et al., 2005).

In the frequency domain, differences in the neural processing of social vs non-social stimuli are typically characterised in adults by increased EEG power in the 5-15Hz range (theta and alpha) (Rousselet et al., 2007). Similarly, older (12-month-old) but not younger (6-month-old) infants also show greater theta power during social compared with non-social viewing contexts (Jones et al., 2006). Greater theta power and changes in alpha power have also been observed in response to child-directed speech and toy play (Orekhova et al., 2006; Stroganova et al., 1998) and joint engagement with an adult (Anderson et al. 2022). Further, making eye contact with an adult before jointly viewing a toy elicits alpha suppression in 9-month-old infants, whilst viewing a toy in the absence of eye contact does not (Hoehl et al., 2014). Just as with developmental changes, though, the question of how fixation-related potentials differ between social and non-social contexts, and how these differences relate to the effects previously observed in the frequency domain, remain unexplored.

To examine these questions, we analysed two datasets of combined EEG and eye-tracking data from 24-month-old children (N=114) and adults (N=108) while they watched stimuli that were either social (an actor singing nursery rhythms) or non-social (dynamic toys). We examined EEG data time-locked to the offset of spontaneous eye movements (i.e., the onsets of fixations) and performed a spectrum of analyses aimed at highlighting the progression of various neural signals across time, frequency, and space (topography). First, we examine increases in fixation related potential (FRP) amplitude over baseline. We consider a range of different aspects of the FRP, including both features specific to FRPs, such as the Spike Potential (SP) (Dimigen et al, 2020), as well as features that FRPs share with passive, task-evoked ERPs, such as the P1, N170, and Nc ERP components. The SP is a maximally frontal negative deflection which ramps up ∼5–10ms before the saccade and reaches its primary peak at saccade onset (Keren et al., 2010). It can also appear time-locked to fixation onsets as a result of micro saccades during a fixation (Dimigen, 2020). The P1 component refers to the first positive deflection in an ERP waveform that typically peaks around 100 milliseconds after the presentation of a visual stimulus and is maximal over medial occipital electrodes (Farroni et al., 2002). The N170 (N290 in infant EEG data) is a negative deflection, peaking at approximately 170ms in adults and 290 ms in infants after stimulus onset and is maximal over lateral-inferior posterior electrodes (de Haan et al., 2003; Guy et al., 2016; Halit et al., 2003). The Nc is observed from approximately 300 to 800 ms without a strictly defined peak latency (for review see de Haan et al., 2003; Reynolds and Richards, 2005, 2009) and is maximal over frontal and central electrodes.

Second, we examine differences in distributions of frequency domain activity across our data, including both during the FRP and times when children are fixating but FRP activity has subsided. Because traditional time-frequency representations essentially measure the summed energy in the signal across the various ERP components, we did this using a more sensitive but less commonly presented output of time-frequency decomposition- the real result of convolution, essentially single trial ERP data that has been band passed filtered at increasing frequency bands from 1-40Hz (Cohen, 2014, p.160). Unlike power, this value can be positive or negative depending on the phase relationship between the signal and the wavelet (see SM Section 3 for more details). We refer to this measure as time-frequency amplitude.

The framework we laid out above generates several research questions with testable predictions:

H1) FRP component activity will differ between children and adults, mirroring what we know about developmental changes in passive ERPs. If this is true, then children should show longer latency P1’s (Taylor et al., 2004). P1 activity should manifest in the frequency domain as theta and alpha in children and adults (Rousselet et al., 2007; Dimigen et al., 2020), and the amplitude of the Nc and N170 should be greater in children than adults (Oades et al., 1997).

H2) Frequency-specific activation associated with FRP components should differ between social and non-social content (i.e., should be context dependent). If true, then we should observe greater frequency-specific increases in spectral amplitude and power (measured in theta and alpha) during the viewing of social vs non-social stimuli, and these differences should be attributable to specific FRP components.

H3) Frequency-specific FRP components will greatly contribute to the frequency domain activation observed in non-event-locked paradigms. If H3 is true we should observe more power during vs after the time course of the FRP, and no differences between total non-event-locked power and power time-locked to FRPs. In order to examine whether frequency-specific FRP components are the result of evoked (additive) brain activity, rather than induced changes in ongoing underlying brain dynamics, we also calculated inter-trial coherence (ITC) (Fig S6) and compared the averaged data with the average after subtraction of the non-phase-locked FRP (Fig S7a).

## 2. Methods

### Ethics statement

Participants were recruited for a longitudinal study running from 2013 to 2019 (STAARS). Informed written consent for the children was provided by the parent(s) prior to the commencement of the study. Ethical approval was granted by the National Research Ethics Service and the Research Ethics Committee of the Department of Psychological Sciences, Birkbeck, University of London. Participant families were reimbursed expenses for travel, subsistence and overnight stay if required. Infants were given a certificate and t-shirt after each visit.

### Participants

For the present analyses data was taken from 114 (51 female), 24-month-old (M = 25.5) typically developing full-term children. Mullen ELC (early learning composite) at 24 months (M = 103.5, SD = 20.6). Social Responsiveness Scale (SRS) raw scores at 36 months (M = 39.9, SD = 31.8). See supplementary materials section 8 for details of exclusion criteria. Analyses presented in the supplementary materials (Figure S1) show that relationships between fixation-related potentials and time-frequency domain activity were highly similar across typical and infants with a family history of a neurodevelopmental condition, meriting their inclusion in this paper.

Adult data was taken from LEAP, a European multisite longitudinal observational study with two complete waves of assessment and an ongoing third assessment; for a comprehensive clinical characterization of the full LEAP cohort, see (Charman et al., 2017). For the present analyses data was taken from 108 participants from the neurotypical control group, ages 18-30 (M = 18.5). No clinical history. See supplementary materials section 8 for details of exclusion criteria.

### Scene-viewing experiment

Participants were presented with two movies of 60-second duration repeated up to three times each per movie (exact numbers are given below). The order of presentation was counterbalanced and interspersed with other elements in the testing battery. Movies were (a) Social: two women telling nursery rhymes with gestures; (b) Non-Social: child-appropriate dynamic toys (e.g. balls dropping down a chute). Similar stimuli have been used in previous studies to examine the effects of socioeconomic status on brain activity (Tomalski et al., 2013) and connectivity in infants with older siblings with ASD (Orekhova et al., 2014).

### Eye tracking data

Stimuli were presented on a 21-inch monitor (refresh: 60 Hz) at a viewing distance of 60 cm. Binocular eye movements were recorded at 120 Hz (300Hz in the adult data) with a Tobii TX300 eyetracker. Eye-tracking data for the social and non-social scene viewing was embedded within recordings from a longer eye-tracking battery. Offline, saccades, and fixations were detected using the I2MC algorithm/toolbox (Hessels et al, 2017). We used the toolbox with its default settings for child and adult data. Synchronisation of the ET and EEG data was done using DIN event markers which were sent in parallel to each data stream.

### EEG data collection

Child EEG data was collected using a 128-channel EGI system; recorded online at 1000Hz, with reference to the vertex. Adult EEG data was collected using a 77-channel Brain Vision system; recorded online at 1000 Hz with reference to the vertex. EEG data was collected from multiple sites. Details of the harmonisaiton across sites are given in supplementary materials section 9.

### EEG data preprocessing

A fully automatic artifact rejection procedure was adopted, following procedures from commonly used toolboxes for EEG pre-processing in adults (Mullen, 2012; Bigdely-Shamlo, et al., 2015) and infants (Gabard-Durnam et al., 2018; Debnath et al., 2020). Full details of the pre-processing procedures can be found in (Haresign et al., 2021). We have used this pipeline in several publications across various developmental datasets and contexts (Marriott Haresign et al., 2023), with only a few minor adjustments to the core settings. In brief, the data were filtered between 1 and 40Hz and re-referenced to a robust average reference. Noisy channels were then interpolated based on correlation: if a channel had a lower than 0.1 correlation to its robust estimate (average of other channels) then it was removed. The mean number of channels interpolated was 4.2 (SD =5.3) for social data and 4.3 (SD =4.4) for non-social data. No channels were interpolated in this way in the adult data. We then removed sections from the continuous data in which the majority of channels contained extremely high-power values. Data were rejected in a sliding 1-second epoch and based on the percentage of channels (set here at 70% of channels) that exceeded 5 standard deviations of the mean EEG power over all channels. For example, if more than 70% of channels in each 1-sec epoch exceeded 5 times the standard deviation of the mean power for all channels then this epoch is marked for rejection. This step is designed to remove data associated with child fussiness and therefore was not applied to the adult data. As a final step to ensure that the EEG data was free from noisy channels (this was particularly important as detailed topographical information was key to our analyses) we ran a second channel interpolation step in which noisy channels that exceed 1000mV were identified and interpolated. For children, this led to an additional 5.7 (SD =7.7) for social movies and 8.4 (SD =19). Overall, 9.9 and 12.7 channels were interpolated on average for social and non-social data, less than 10% of the total number of channels for both conditions. For adults less than 1 channel on average was removed during this step (M = 0.81 for social, M = 0.89 for non-social). We also conducted additional ICA cleaning, but our analyses presented in the SM (see Figures S2 and S3) suggested that this did not successfully attenuate artifact in the child data and attenuated genuine neural activity in adults. Therefore, we elected to carry out the current analyses on data not cleaned using ICA.

### Combined Eye-tracking and EEG datasets

*Children*. Of the 114 possible eye-tracking (ET) datasets,10 had missing data for the social and non-social movie viewing conditions and 2 recordings ended prematurely resulting in an insufficient amount of data. Of the 102 possible ET datasets, each had 2.9 (out of 3 possible) on average trials of each movie viewing, totalling 296 trials of movie presentations. Of these, the I2MC algorithm identified at least one fixation in 241 trials (∼19% loss of data). 9 EEG datasets had too few data samples to preprocess (∼<2s of data) (9% loss of data). Of these, 81 (87%) of the 93 EEG datasets had paired ET data.

*Adults*. Of the 48 possible eye-tracking (ET) datasets, 0 were missing data. Each had 2 on average trials of each movie viewing, 96 possible ET trials. Of these, the I2MC algorithm identified at least one fixation in 86 trials (∼11% loss of data). Of the 108 possible EEG datasets 2 were missing data for the social and non-social movie viewing conditions. 5 EEG datasets had too few data samples to preprocess (∼<2s of data) (5% loss of data). Of these 48 (48%) of the 101 EEG datasets had paired ET data.

Our analyses time-locked brain changes to the offsets of eye movements. Of note, we decided not to exclude trials in which subsequent eye movements occurred during the time window we were analysing, because our descriptive statistics showed that subsequent saccades were evenly distributed in time, and because of the close correspondence already observed between active fixation-related potentials obtained using these methods and passive, stimulus-evoked ERPs (described below).

### Time-frequency analysis

Time-frequency representations of ERP amplitude and power often show broadband increases (Rousselet et al., 2007), that span the entire time range of the ERP waveform. Thus, in essence, they measure the summed energy in the signal across the various ERP components. This approach can, however, be insensitive to how different ERP components are differentially contributing to increases in spectral power. In this paper, we use a less commonly presented output of time-frequency decomposition-the real result of convolution, essentially single trial ERP data that has been band passed filtered at increasing frequency bands from 1-40Hz (Cohen, 2014, p.160). This was done using a complex Morlet wavelet convolution using wavelets that increased in 18 logarithmically spaced steps and the number of cycles increased from 3-10 cycles logarithmically. Unlike power, this value can be positive or negative depending on the phase relationship between the signal and the wavelet. In the supplementary materials section 3, we present more details on how this approach allowed us to better understand the contributions of different components of ERPs/FRPs.

Electrode groupings used for all analyses are given in SM (Figure S4).

## 5. Results

Our overall hypothesis was that neural activity time-locked to eye movements generates frequency-specific neural activity that supports children’s visual attention and cognition. To examine this, we compared neural activity in child and adult data associated with different contexts – i.e., the processing of social vs non-social stimuli. Our aims were: i) to explore whether any observations of time-frequency FRP activity reflected context-independent or context-specific processes and ii) to investigate similarities between fixation-related activity and previously documented effects of age and stimulus type in the frequency domain.

### 3.1 Differences in saccadic/ fixation behaviour

Figure 1 provides descriptive statistics of children and adult eye movements while viewing social and non-social stimuli, and a descriptive illustration of a fixation-related potential.

**Figure 1.**
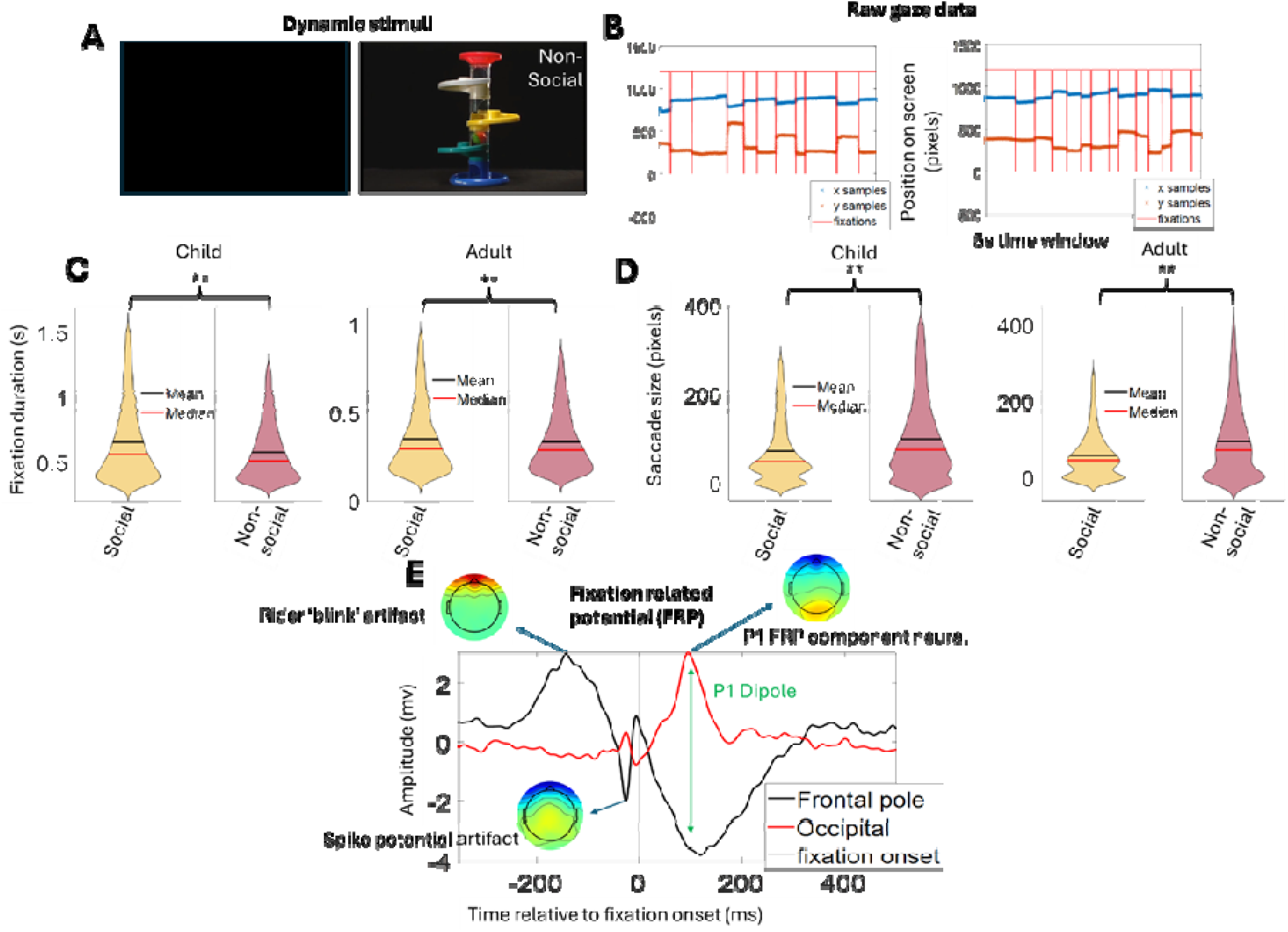
**illustration of social and non-social condition, eye movement differences and fixation-related potential.** A. Sample screenshots from the dynamic social and non-social stimuli. B. Raw eye tracking gaze data child and adult – shows identification of fixations. C. Fixation durations for child and adult social and non-social stimuli. ** indicate the significance of p < 0.01 based on two-sample t-tests. D. Saccade sizes for child and adult social and non-social stimuli. ** indicate the significance of p < 0.01 based on two-sample t-tests. E. EEG data was time-locked to the onset of fixations, and we examined neural and artifactual EEG signals that happen as a result of a new fixation.

We first investigated differences in children’s and adults’ eye movements while viewing social and non-social stimuli, considering both the average rate of sampling (inverse to fixation duration) and the average size of saccades across both contexts. Results of the two-sample t-tests suggested that children t(21,184) = 19.1, p < 0.01 and adults t(17,213) = 7.5, p < 0.01 both fixated longer on average during the social vs non-social condition and made smaller saccades during social vs non-social viewing, children t(21,184) = −12.7, p < 0.01, adults t(16,993) = −14.0, p < 0.01. Thus, suggesting that the content-specific differences in visual foraging and processing might arise both at the visual motor and perceptual level. At the neural level (FRPs) there is a clearly observable mix of artefactual signals generated from the eye movement and neural processing (Figure 1C).

### 3.2. Differences in FRP components between children and adults

In order to examine differences between children and adults’ cortical activity relative to eye movements (our first hypothesis, H1), we compared peak latencies and amplitudes of FRPs across four key components based on previous literature (Dimigen et al., 2011; Dimigen, 2020): SP (artifact), P1 (neural), Nc (neural), N170 (neural) (Fig. 2A, Fig. 3). With the exception of the SP, all of these FRP components each have an analogous passive ERP component (although the Nc is observed in children but not adults (Richards, 2010)). Here, we focused on a limited range of statistical comparisons – comparing peak latencies and amplitudes over set time windows relative to fixation onset (SP, −100 to 50ms, P1, 0-200ms, Nc, 300-600ms and N170, 100-250ms), chosen based on previous literature and our own observation of the data, using two-sample t-tests. Results of the two-sample t-tests (visualised in Fig. 3) indicated several differences in FRP waveforms between children and adults, namely: significantly longer P1 t(283) = 2.8, p < 0.01 (Fig. 3D) and SP t(283) = −4.8, p < 0.01 latencies (Fig. 3A) (measured from fixation onset). In children, the peak of the SP occurred earlier pre-fixation and the peak of the P1 happened later post-fixation onset.

**Figure 2.**
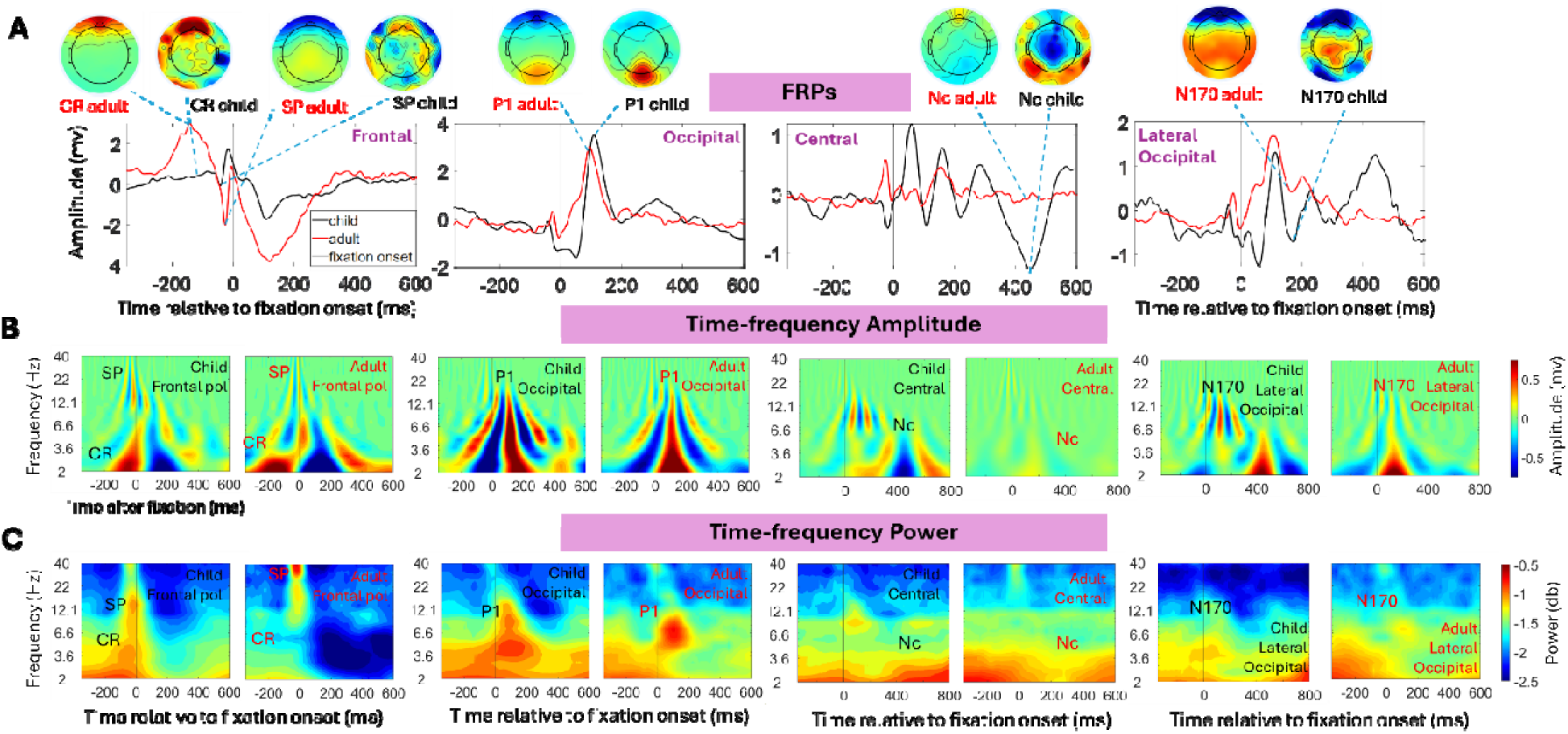
**Neural dynamics of spontaneous fixations in children and adults.** A. Fixation-related potentials (FRPs) from different electrode groupings, differentiating frontal pole, occipital, central, and lateral occipital. Topoplots show, for key components, the topographical distribution of activity at the peak of the labelled potential. CR – corneo-retinal dipole; SP – spike potential; P1 – P100 ERP component; Nc – Negative Central; N170 – N170 ERP component. Time 0 indicates the onset of a fixation. B. Time-frequency amplitude plots for the same FRPs. C. Time-frequency power plots for the same FRPs. Figure 3 shows violin plots and statistical analyses of the results shown in Figure 2.

**Figure 3.**
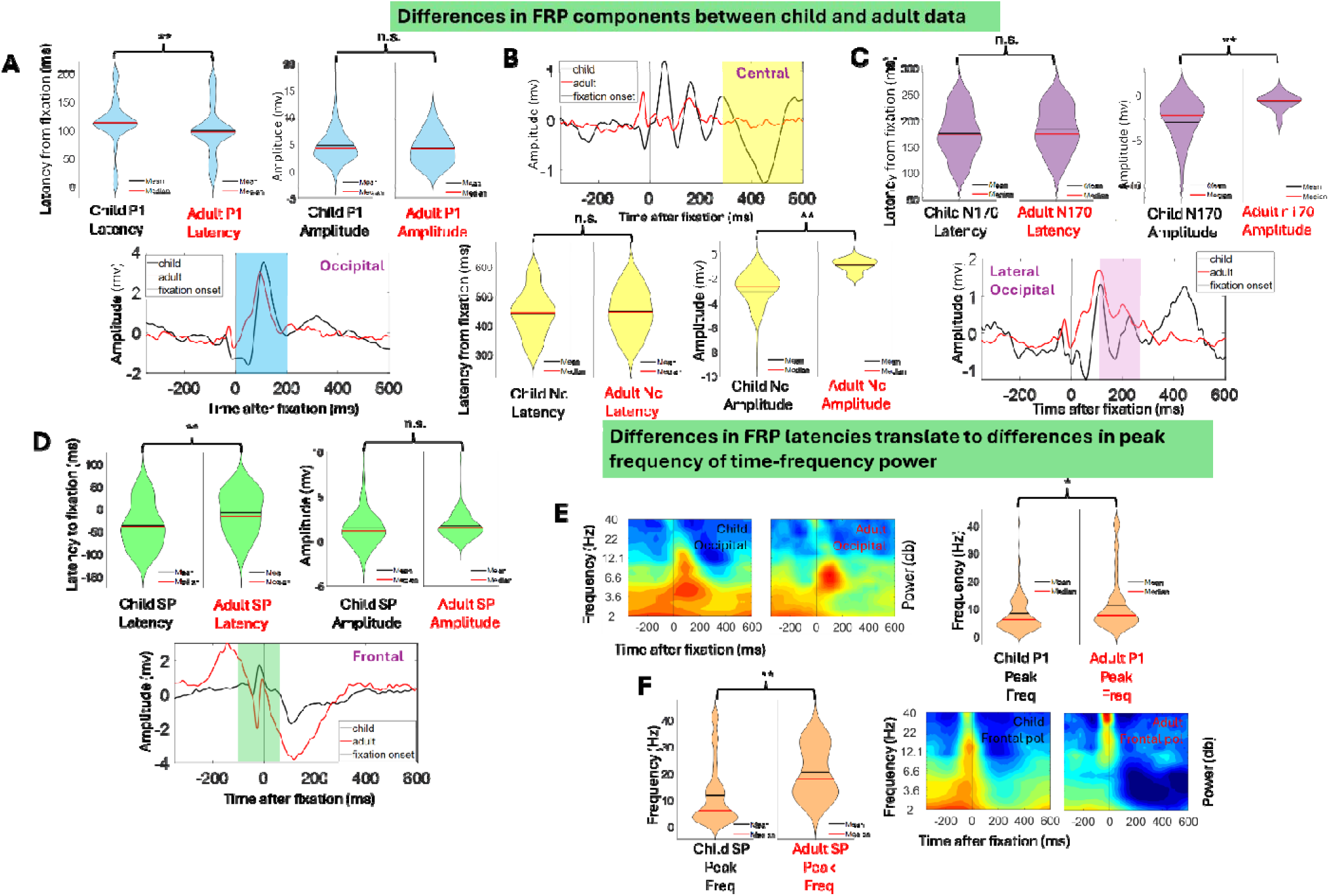
**Differences in latency, amplitude and peak frequency power between child and adult FRP components.** For A-D top plots show distributions, mean, and median of peak activity highlighted in corresponding coloured time windows of FRP plots (bottom plots of A-D) for child and adult data. The same time windows were used for time (FRP) and time-frequency analyses (see section 3.2 for more details). ** indicate the significance of p < 0.01 based on two-sample t-tests. ‘n.s’ refers to tests that were non-significant. A. Highlights (in blue) comparisons of latency and amplitude between child and adult P1 FRP components. B. Highlights (in yellow) comparisons of latency and amplitude between child and adult Nc FRP components. As expected based on previous literature (Richards, 2010) the Nc is present in children but absent in adults. C. Highlights (in purple) comparisons of latency and amplitude between child and adult N170 FRP components. D. Highlights (in green) comparisons of latency and amplitude between child and adult P1 FRP components. E. Shows child and adult occipital FRP power relative to fixation onset and distributions, mean and median of peak power for P1 FRP component activity highlighted in the blue section of Fig. 3A. * indicates the significance of p < 0.05 based on two-sample t-tests. F. Shows child and adult frontal pole FRP power relative to fixation onset and distributions, mean and median of peak power for SP FRP component activity highlighted in the green section of Fig. 3D. ** indicate the significance of p < 0.01 based on two-sample t-tests.

To examine how these differences in children’s and adults’ processing were manifesting cortical oscillatory activity (H1), we next sought to examine time frequency amplitude (Fig. 2B) and power (Fig. 2C) relative to the offsets of eye movements. These analyses indicated that eye movement artifact and neural processing (P1) are clearly differentiable in time and frequency space (see SM section 3). Importantly, however, these analyses revealed that several differences between children and adults FRP latency manifested as differences in frequency band oscillations. Saccade-related activity manifests as a brief increase in gamma time-locked to the saccade in adults (above 10Hz, M = 30.4); the same saccade-related activity manifests as beta in children (above 10Hz, M = 14.1) (Fig 2C). Results of two sample t-tests revealed differences in mean peak frequency of SP for children vs adults t(283) = − 2.9, p < 0.01 (Fig. 3F). P1-related FRPs– manifest as theta and alpha activity (M = 10.7) in adults and children (M = 8.2) but at a lower frequency in children t(283) = −2.4, p = 0.02 (Fig. 3E). Therefore, differences in FRP latencies between children and adults manifest as differences in peak frequency. These findings for active FRPs mirror what we already know of developmental trends of these components for passive ERPs. Crucially however these differences also manifest as differences in the time-frequency expression of these signals.

### 3.3. FRP components manifest as frequency-specific neural oscillations in children

In section 3.2 we introduced the possibility that differences in FRP component latency (H1) between children and adults were manifesting as differences in peak frequency of oscillatory power. Here, in this section, we sought to further quantify where FRP component activity was resulting in frequency-specific neural activity in children and adults (H1). To do this we examined whether neural activity during the FRP increased against baseline (pre-fixation onset) and whether any increases were confined to neural oscillations at specific frequencies. The time windows used in these analyses are highlighted in the FRP plots (Fig 4A, 4B, 4C, left plots). To estimate significance, we used one-dimension cluster-based permutation statistics in frequency space, against an alpha value of 0.01.

**Figure 4.**
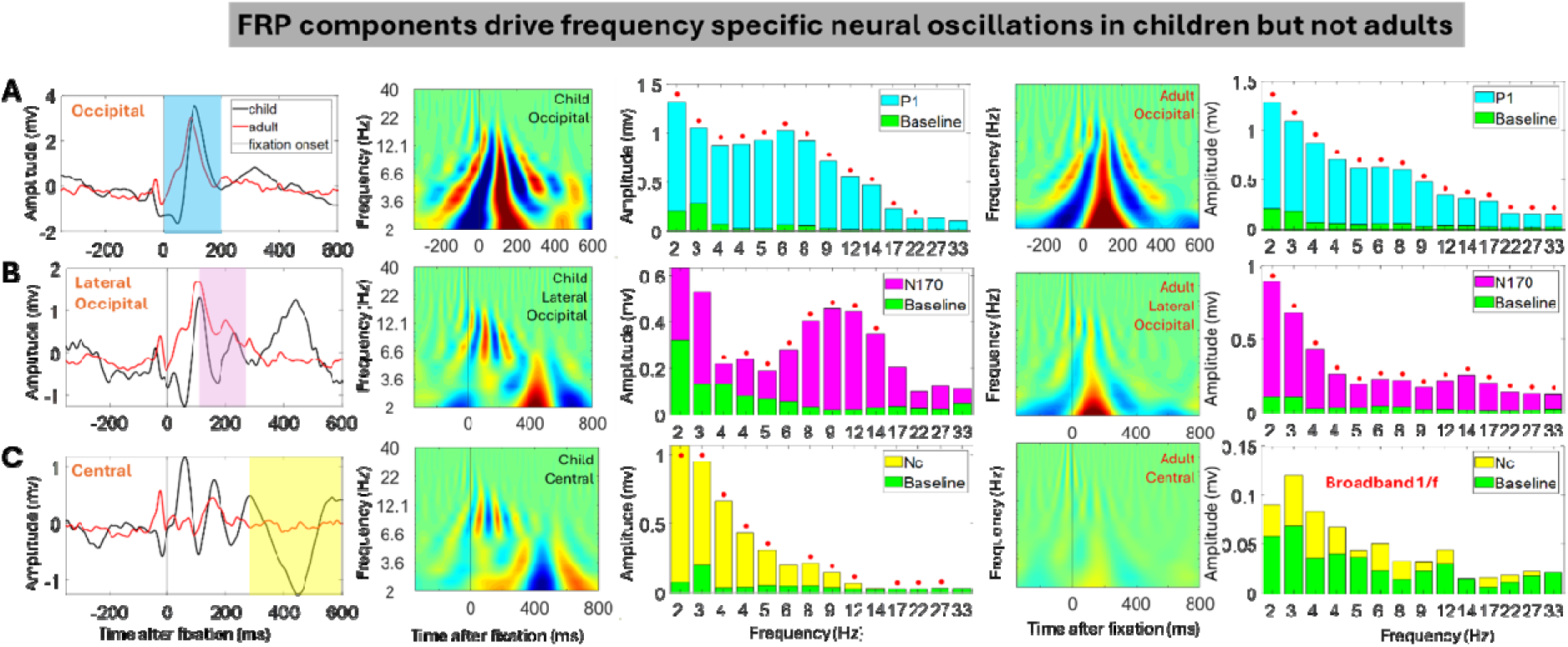
**Time-frequency amplitude changes pre vs post fixation onset in children and adults.** Time windows used for feature extraction are highlighted in the FRP plots. The same time windows were used for time (FRP) and time-frequency analyses (see section 3.2 for more details). A. FRP P1 activity (section in blue) in children and adults. Corresponding time frequency amplitude plots (row A plots 2 and 4) and histograms comparing time-frequency amplitude pre-vs post-fixation onset (for section highlighted in blue) for both children (row A plot 3) and adults (row A plot 5). B. FRP N170 activity (section in purple) in children and adults. Corresponding time frequency amplitude plots (row B plots 2 and 4) and histograms comparing time-frequency amplitude pre-vs post-fixation onset (for section highlighted in pink) for both children (row B plot 3) and adults (row B plot 5). C. FRP N170 activity (section in yellow) in children and adults. Corresponding time frequency amplitude plots (row C plots 2 and 4) and histograms comparing time-frequency amplitude pre-vs post-fixation onset (for section highlighted in pink) for both children (row C plot 3) and adults (row C plot 5).

For adults, P1 and N170 FRP components were associated with increases in amplitude over all frequencies compared to baseline. As expected, no increases were associated with the Nc FRP component. This suggests that, for adults, FRPs generate a relatively broadband response. For children, the N170 FRP component was associated with increases in amplitude in the frequency range of 3-14 Hz only. The P1 component was associated with increases in amplitude in the frequency range 2-22 Hz. Lastly, the Nc component was associated with multiple separate clusters of increased amplitude; cluster 1 frequency ranges 2-5 Hz, cluster 2 frequency range 7-11 Hz (which is likely driven by dipolar activity associated with the N170), cluster 3 17-26 Hz. This suggests that in children FRPs are associated with frequency-specific changes in activity.

### 3.4. Frequency-specific neural activity differs during social and non-social contexts, consistent with changes in global oscillatory activity

Having established that certain FRP components manifest as frequency-specific cortical oscillations we examined whether this activity was context-dependent (H2). We were also interested in the relationship between context-specific effects at the FRP level (i.e., transient cortical excitability at the onset of fixation) and neural oscillations documented using non-event-locked analyses (H3).

First, to address H2, we examined differences in theta band power between social and non-social movie viewing for adults (4-8Hz) and children (2-7Hz). We examined differences across the EEG data globally (i.e., before/ not event-locked to fixation onsets) and in the event-locked data. We first looked at this topographically, using 2-dimensional cluster-based permutations (with an alpha value of 0.05) on topographical illustrations of social and non-social theta power (Fig. 5A). The results of these analyses revealed significant clusters of increased theta power for social vs non-social movie viewing in children, but not adults, both at the global and the event-related level. These clusters were largely localized occipital and frontocentral electrodes. No significant differences were found between social and non-social movie viewing in adults (Fig. 5D-F). As we observed in previous sections, key FRP components manifest as frequency-specific neural oscillations: P1 associates with theta and alpha and the N170 associates with alpha. We next looked at whether there are differences between social and non-social in the time-frequency manifestations of key FRP components. The results of these analyses revealed two significant clusters of increased amplitude – which were associated with the time-frequency manifestations of the P1 and N170 (Fig. 5B). Further we observed that the Nc manifests as low theta and shows greater (more negative) amplitude for non-social vs social conditions over central electrodes (Fig. 5C). Again, adults showed no significant differences in time frequency amplitudes for social vs non-social FRPs. These results suggest that previously observed differences in oscillatory activity between viewing contexts may be explained via changes in cortical excitability time-locked to eye movements.

**Figure 5.**
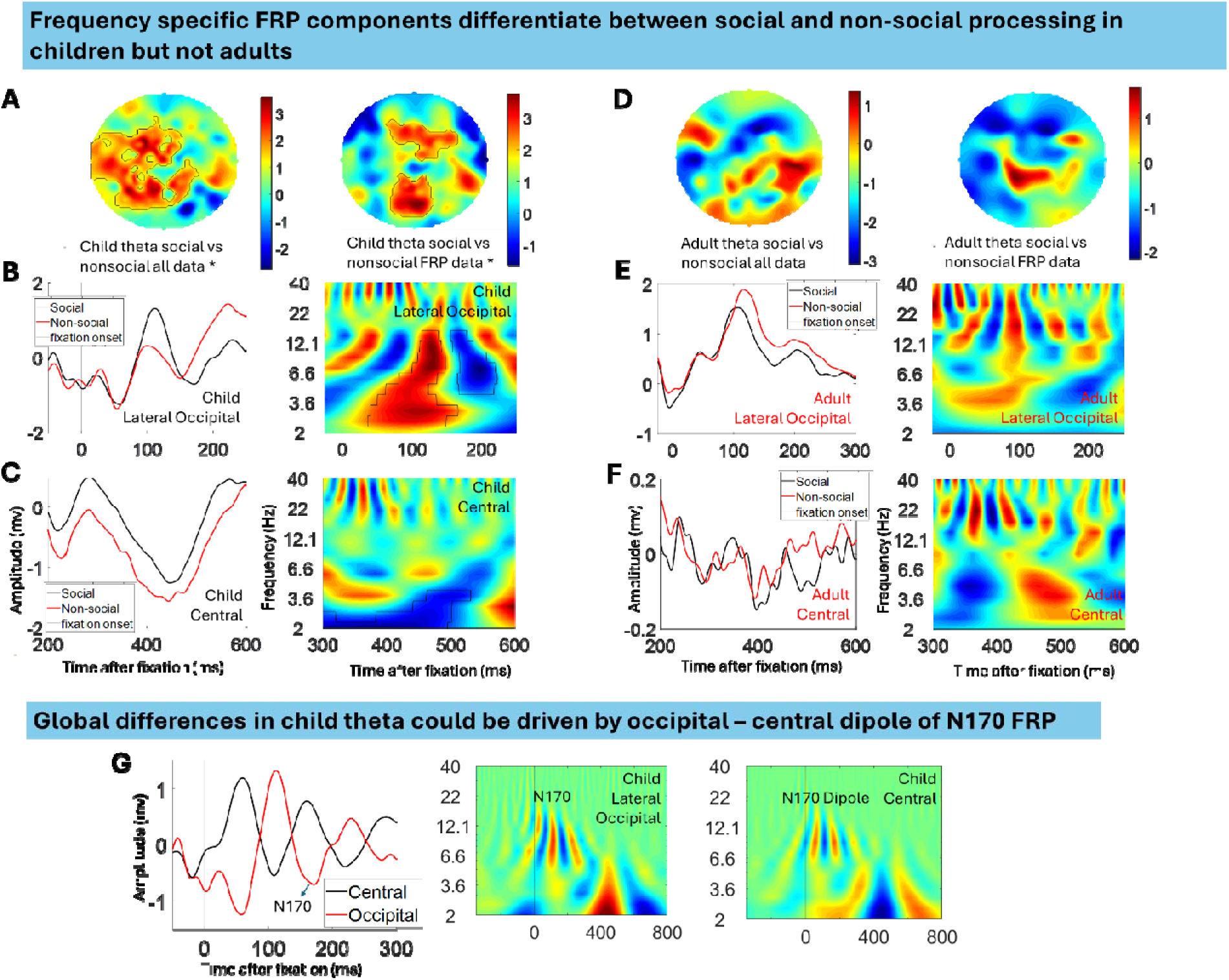
**Differences in frequency-specific neural activity of FRP components between social and non-social contexts in children and adults. For all plots black outlines show significant clusters of increased power (p<0.05).** A. Global and event-locked topographical differences in child theta (2-7Hz) social vs non-social. B. Differences between child occipital N170 social vs non-social. Differences in N170 manifest as differences in time frequency theta and alpha. C. Differences between child central Nc social vs non-social. Differences in Nc manifest as differences in time frequency theta. D. Global and event-locked topographical differences in adult theta (4-8Hz) social vs non-social. E. Differences between adult occipital N170 social vs non-social. F. Differences between adult central Nc social vs non-social. G. Clusters of increased theta power could be driven by N170 dipole as significant clusters of increased theta power are driven by two topographical sources in children (Fig. 5A) an occipital and a fronto-central cluster. For all plots, black outlines show significant clusters of increased power (P<0.05).

To address our third hypothesis (H3), we first explored differences between the topography of child theta for all data, during an FRP (defined as the first 850ms following the onset of each fixation), and post-FRP (defined as the period from 850ms until the end of the fixation) (Figure 6). We identified focal frontal clusters of theta during the global non-event-locked data (Figure 6A), and frontal and occipital clusters during the FRP (Figure 6B) that were absent outside of the FRP (Figure 6C). When we statistically compared topographical images using cluster-based permutation statistics we found that theta levels during the FRP were significantly higher (Figure 6D) than all data. We also identified a marginally non-significant pattern that theta power was higher during an FRP level vs outside of the FRP (Fig. 6E). Figure S5 shows the same pattern present in adults.

**Figure 6.**
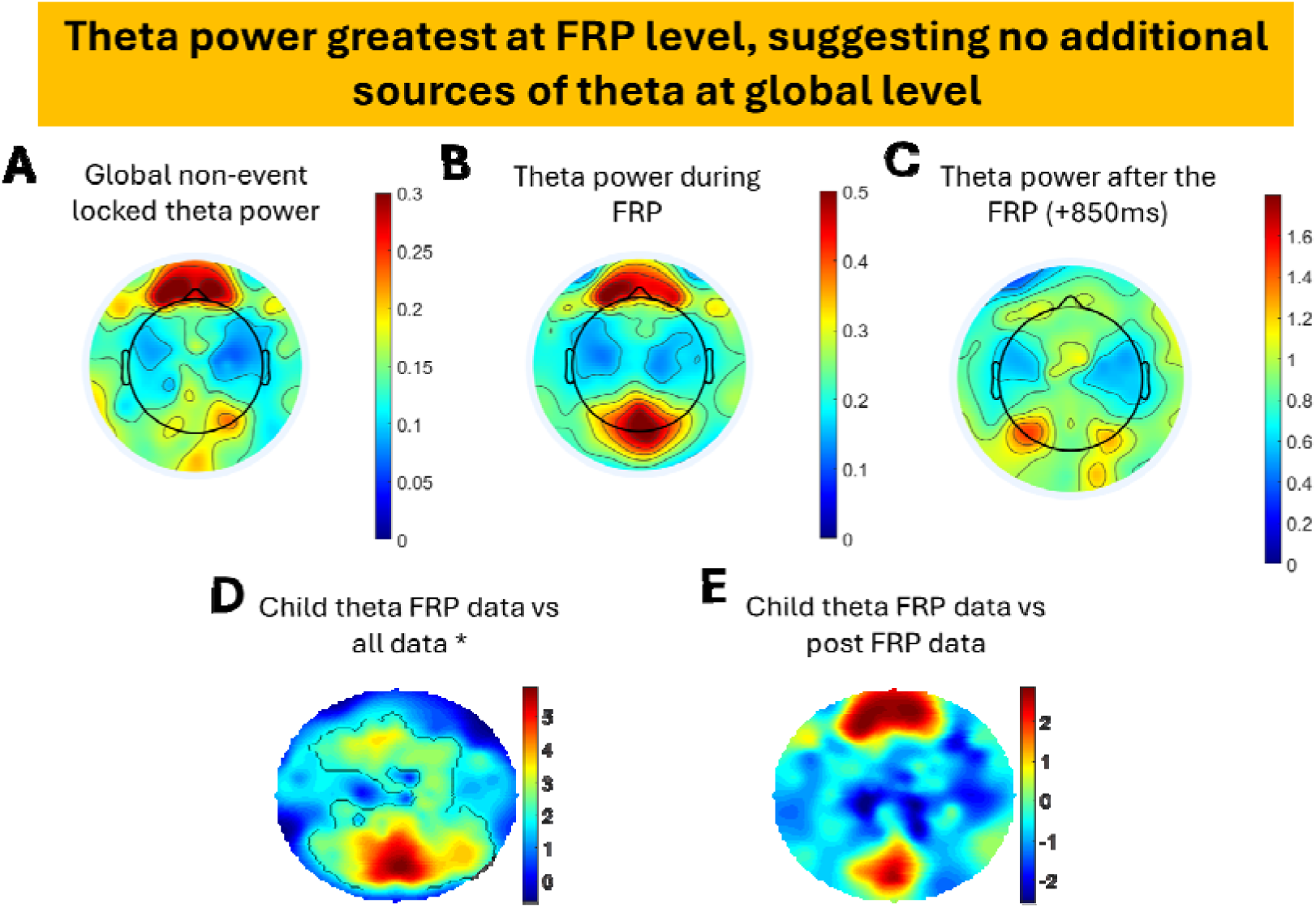
**Child theta (2-7Hz) differences across all data, FRP data and post FRP data. Equivalent plots for adult data are shown in Fig S5. For all plots black outlines show significant clusters of increased power (p<0.05).** A. Topographical distribution of child theta power across entire data sample, not time-locked to fixation onsets. B. Topographical distribution of child theta power during fixation-related potential (FRP) time-locked to the onset of fixation. C. Topographical distribution of child theta power post FRP (+850ms after fixation onset) D. Theta power during data segments where FRPs are present vs all data. E. Theta power during data segments where FRPs are present vs not present. The figure shows theta is largely driven by the activity of FRP as no sources of increased theta at the global level (all data) and outside of the FRP. See Figure S5 for adult data. For D and E black outlines indicate significant clusters.

Finally, to test whether changes in oscillatory activity that occur time-locked to fixation onsets are the result of evoked brain responses, rather than induced changes in ongoing underlying brain dynamics, we conducted two additional analyses. First, we calculated changes in inter-trial coherence (ITC) for child FRP analyses of occipital and frontal pole electrodes and compared them with the time-frequency amplitude and power changes already shown in Figure 2 (Fig S6). Changes in ITC were observed following fixation onset, but these closely mapped the observed power changes, showing the difficulty of disentangling power and phase estimate (Burgess, 2013 van Diepen & Mazaheri, 2018). Second, we calculated child FRP theta power based both on the entire averaged data and on the averaged data after the subtraction of the non-phase-locked FRP (Fig S7A). Oscillatory activity was strongly attenuated after subtraction of the FRP suggesting that changes in oscillatory activity are the result of fixation-evoked brain events.

Finally, we re-examined previous analyses (Perapoch Amado et al., 2023; Phillips et al., 2024; Wass et al., 2009) which identified time-locked associations between frontocentral theta power and infants’ attention to objects and people around them, which have been used to motivate claims that theta oscillations index active learning in infancy (Begus & Bonawitz, 2020). Results show that the removal of phase-locked FRP completely attenuates the cross-correlations previously observed between theta power and visual attention. Overall, these analyses are suggestive that topographical and temporal variance in theta power is most notably contributed to by dynamics associated with FRP-level activity.

## 6. Discussion

We examined changes in brain activity time-locked to spontaneous eye movement offsets using combined EEG and eye-tracking data from 24-month-old children (N=114) and adults (N=108). Participants viewed either social stimuli (an actor singing nursery rhymes) or non-social stimuli (dynamic toys). EEG data were analysed to track neural signal progression across time, frequency, and topography. Our aim was to determine how transient changes in cortical excitability linked to eye movements shape oscillatory brain activity and how these effects vary with development and viewing context.

First, we investigated how differences between children and adult’s fixation-related potential (FRP) components manifested in time-frequency space. We hypothesised (H1) that FRP component activity would differ across age groups, reflecting developmental changes observed in passive, task-evoked ERPs. Prior studies indicate that task-evoked visual ERPs evolve with age (Taylor et al., 2001; Itier & Taylor, 2004a,b; Batty & Taylor, 2006). Research on facial ERPs in children has shown age-related decreases in P1 and N170 amplitude and latency (Taylor et al., 2004; see also Buchsbaum et al., 1974; Brecelj et al., 2002). Our findings for active FRPs align with these trends: children’s P1 latencies were longer, and their N170 amplitudes were higher (Figure 3), suggesting that early-stage visual processing follows the same developmental trajectory whether information is actively selected or passively received. Visual processing becomes faster and more efficient over development, requiring fewer neural resources to process equivalent stimuli.

Next, we examined whether FRP components time-locked to eye movements manifest as frequency-specific neural oscillations. We compared time-frequency amplitude changes before (baseline) and after eye movement onset (Figures 2-4). We hypothesised that FRP components would induce frequency-specific activity beyond the broadband activation reported in previous literature. In adults, P1 and N170 FRP components were associated with amplitude increases across all frequencies (Figure 4), with no increases linked to the Nc FRP component. This finding aligns with prior research (Dimigen et al., 2011; Leszczynski & Schroeder, 2019; Maldonado et al., 2008; Rajkai et al., 2008; Rousselet et al., 2007), indicating that adult FRPs generate a broadband response without driving frequency-specific activity. In contrast, children’s FRPs displayed more differentiated patterns (Figure 4). While P1 showed broadband amplitude increases (2-22 Hz), the N170 manifested as alpha-range activity (6-12 Hz, Figure 4B), and the Nc appeared predominantly in the theta range (2-5 Hz, Figure 4C). These results suggest that FRPs contribute to oscillatory activity differently across age groups.

We also examined whether these frequency-specific FRP components showed content specificity. We predicted (H2) that frequency-specific FRP activation would differ between social and non-social stimuli. To test this, we first looked at differences in neural activity between social and non-social content for adults and children (Figure 5). Motivated by previous literature (Jones et al., 2006; Orekhova et al., 2006; Rousselet et al., 2007) we concentrated on the theta band. Our 2D cluster-based topographical analysis (Figure 5A) revealed increased theta for social versus non-social stimuli across occipital and fronto-central electrodes in children, but not in adults, supporting earlier findings (Jones et al., 2006; Orekhova et al., 2006). Time-frequency FRP analyses (Figure 5B) identified two significant clusters corresponding to P1 (Figure 4A) and N170 (Figure 4B), consistent with the task-evoked ERP literature (Conte et al., 2020; Xie & Richards, 2016). Nc amplitudes were more negative for non-social conditions over central electrodes (Figure 5C), consistent with previous research (Gui et al., 2021). These results suggest that social and non-social stimuli elicit distinct oscillatory responses in children, with theta activity particularly influenced by eye movement-locked cortical excitability.

These findings parallel language research, where the N400 ERP component—linked to semantic processing (Kutas & Federmeier, 2011; Neville et al., 1992; King & Kutas, 1995)— has been found to overlap with increases in theta power (4-8 Hz) associated with semantic integration (Bastiaansen et al., 2002, 2010; Davidson & Indefrey, 2007; Hagoort et al., 2004; Hald et al., 2006; Maguire et al., 2010; Wang et al., 2012). Our results extend this work by demonstrating: (i) context-, topography-, and frequency-specific associations between transient brain responses and oscillatory activity that align with developmental trends; (ii) parallels between passive, task-evoked ERPs and active, eye movement-locked paradigms; and (iii) evidence that transient responses linked to eye movements may explain prior observations from non-event-locked paradigms.

Finally, we tested our prediction (H3) that frequency-specific FRP components contribute to frequency-domain activation seen in non-event-locked paradigms. We focused on theta activity, considered a marker of individual differences in young children (Begus & Bonawitz, 2010; Braithwaite et al., 2020; Jones et al., 2020; Wass et al., 2019). Theta power increased during an FRP (Figure 6B), consistent with sources originating from P1, Nc, and N170 FRP components (Figures 2 & 5G). Crucially, however, Figure 6C shows no increased theta at the global level (all data) outside the FRP window (Figures 6D-E).

Contrary to prior reports, we also found no evidence that saccade timings were phase-locked to ongoing oscillatory activity (Figure S6). Given the small effect sizes in previous studies (e.g., ΔPLI of 0.02 using MEG (Pan et al., 2021) and 0.01 in macaque frontal eye field recordings (Shaverdi et al., 2023)), our analyses may have lacked sensitivity to detect such subtle effects.

Overall, our results suggest that frequency-specific neural patterns which occur time-locked to eye movements are both age- and context-dependent. In children, these relationships were more pronounced between different viewing contexts (Figures 3-4). Our findings have two key implications. First, they suggest that developmental changes observed in frequency-domain analyses may be better understood as fine-grained, movement-induced brain states. Our analyses allow us to clearly differentiate between eye movement artifact and genuine changes in neural activity that occur time-locked to eye movements (Figure 2). They suggest that transient increases in genuine oscillatory neural activity (e.g., transient increases in theta both directly via the Nc and via dipoles associated with fixation-related P1 and N170 components) occur in the time period following the onset of a new fixation. These build on previous suggestions that eye movements, which naturally occur at about 3 times per second, may generate transient broadband increases in cortical excitability in the theta range. They are also distinct from previous observations that eye movements may be timed to co-occur with the phase of underlying oscillatory activity – which, as discussed above, we did not find any evidence of here. Our present results build on these previous observations in suggesting that micro-movements play a crucial role in shaping neural dynamics.

Methodologically, our study also carries significant implications. Our findings call into question past studies that examined neural activity without measuring fine-grained eye movements. These include studies that measured individual differences in non-event-locked brain activity (e.g., Begum-Ali et al., 2022; Marshall et al., 2002; Jones et al., 2020; Tomalski et al., 2013; Troller-Renfree et al., 2022) and those that explore developmental changes (Cellier et al., 2021; Cui et al., 2020; Marek et al., 2015; Marshall et al., 2002; Stanyard et al., 2024; Voytek & Kramer, 2015; Wilkinson et al., 2024). They also include those that observed changes in brain activity while performing cognitive tasks (Begus et al., 2015; Bell, 2001; Hendry et al., 2025; Orekhova et al., 2006; Xie et al., 2018). Our findings suggest, in studies that do not track fine-grained eye movements, frequency-specific differences in neural activity observed may be attributable to differences in the timing, frequency, or size of eye movements. Although it remains possible that non-event-locked differences in the frequency domain could be driven by brain differences in the absence of behaviour differences (e.g., differences in the amplitude/power of FRPs, even when identical eye movements are taking place) our results suggest that these effects are likely to be smaller than eye-movement-related differences.

To illustrate this, we re-examined previous analyses (Perapoch Amadó et al., 2023; Phillips et al., 2024; Wass et al., 2018) which observed time-locked increases in theta power during attentional engagement. These have been used to motivate theoretical claims that (for example) theta oscillations index active learning in infancy (Begus & Bonawitz, 2020). When we repeated the analysis whilst tracking fine-grained eye movements, which allowed us to calculate and subtract the average FRP (Figure S6E), we found that previously observed time-locked associations are entirely removed. Future work should also examine whether other within- and between-individual differences in brain activity are attributable to behavioural differences in the same way.

Previous authors have noted that: “[i]t remains a possibility that changes in low-frequency neural dynamics currently attributed to particular cognitive functions are confounded with changes in spatio-temporal patterns of active visual exploration” (Leszczynski & Schroeder, 2019). Our present findings are consistent with this. Rather than suggesting that variance in global task-modulated oscillatory activity is a result of the brain’s passive and reflexive information processing, they emphasise the importance of active, transient, movement-induced brain states. Understanding how fine-grained neural activity changes relative to eye movements seems important, then, for understanding diverse known features of brain development, such as the shift in peak frequency from ∼5Hz to ∼8Hz alpha peak with age (Cellier et al., 2021; Wilkinson et al., 2024); the decrease in the aperiodic slope of brain activity (Cellier et al., 2021; Schaworonkow et al., 2021; Stanyward et al., 2024; Wilkinson et al., 2024), and how the role of global task-modulated oscillatory activity in reflexive information processing changes over development.

