## Supplemental materials for "Cortical Excitability during Fixations Drives Frequency-Specific Neural Activity in Children and Adults"

1. **Comparison of FRP activity in typically developing, at risk ASD and ADHD group data**

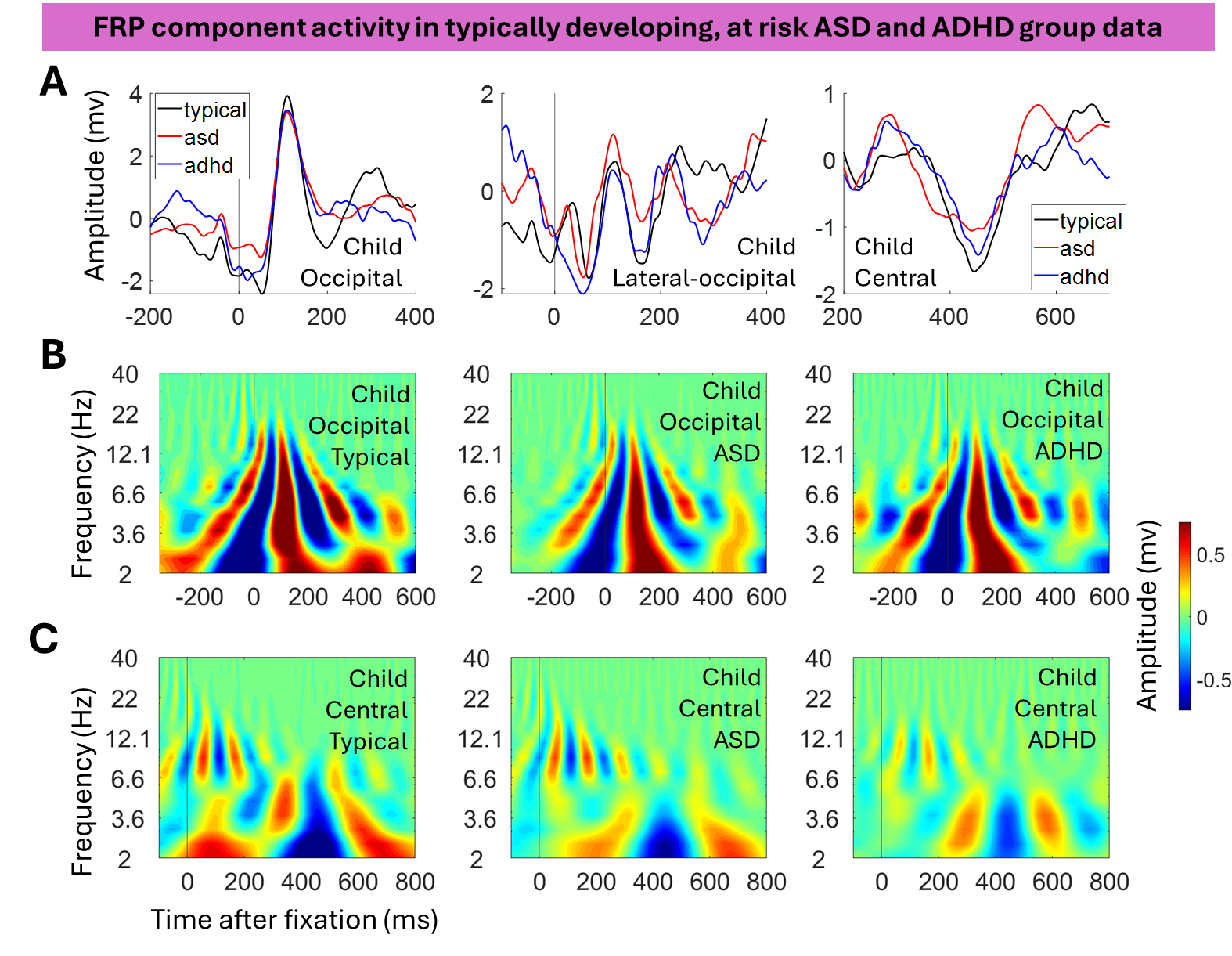

**Figure S1. Neural dynamics of spontaneous fixations in typically developing children and children at risk of ASD and ADHD.**

A. Fixation-related potentials (FRPs) from different electrode groupings, differentiating frontal pole, occipital, central, and lateral occipital. Plots show FRP from three groups typically developing, at-risk ASD, and at-risk ADHD.
B. Time-frequency amplitude plots for the same FRPs over occipital electrodes. Plots show FRP from three groups typically developing, at-risk ASD, and at-risk ADHD.
C. Time-frequency amplitude plots for the same FRPs over central electrodes. Plots show FRP from three groups typically developing, at-risk ASD and at-risk ADHD.

Figure S1 examines FRP activity in typically developing, at-risk ASD and ADHD group data. It can be seen that the same components are present in all groups (panel A) and these components are associated with similar patterns of oscillatory activity (panels B and C). Most crucially, the time-frequency amplitude plots for both occipital activity (Fig S1B) and central activity (Fig S1C) show highly similar patterns of association between fixation-related potentials and time-frequency analyses, such that including the at-risk ASD and ADHD group data alongside typically developing group data if anything is a source of type 2, not type 1 error. In a separate forthcoming paper, we shall specifically compare differences in time-frequency domain activity time-locked to fixations; however, this is a separate research question to the present paper.

2. **Differences in FRP activity before and after ICA cleaning**

Building on previous work (Marriott Haresign et al., 2021), we experimented with using ICA following the most up-to-date procedures for dealing with EEG data contaminated with eye movement artifact (Dimigen, 2020) (we refer the reader there for procedural details) (see Figure S2). It is important to note that we did not follow Dimigen’s procedures exactly but rather used a modified version. We did not use the information from the eye tracker to identify micro saccades and correct for micro saccadic induced artifact.

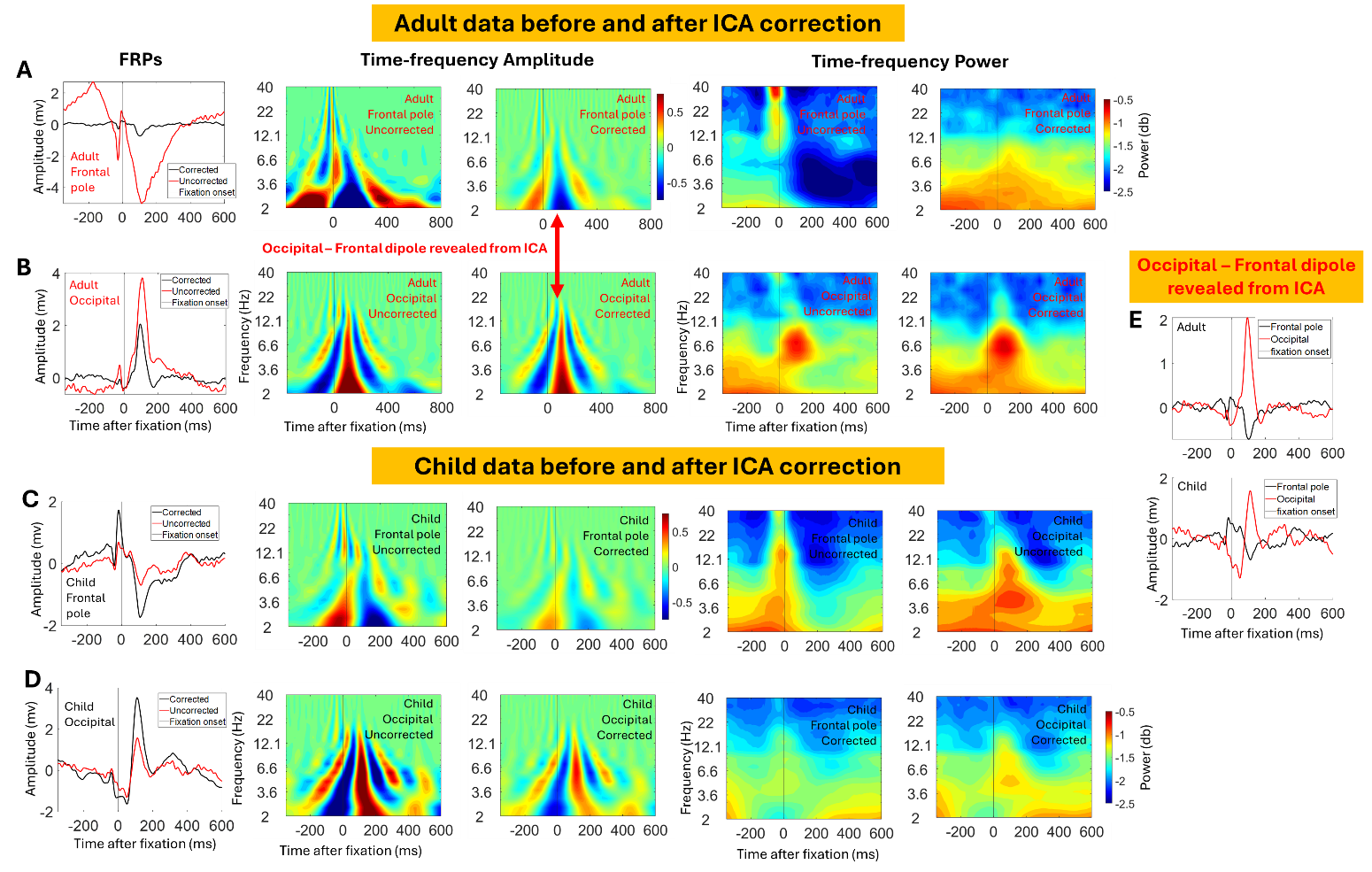

**Figure S2. Results of correction of eye movement artifact using overweighted ICA in children and adults.**

A. Adult FRPs, time-frequency amplitude and power over frontal and occipital electrodes before ICA correction.
B. Adult FRPs, time-frequency amplitude and power over frontal and occipital electrodes after ICA correction.
C. Child FRPs, time-frequency amplitude and power over frontal and occipital electrodes before ICA correction.
D. Child FRPs, time-frequency amplitude and power over frontal and occipital electrodes after ICA correction.
E. Dipole between occipital and frontal electrodes after ICA correction in adult and child data.

To evaluate the effect of the cleaning, we compared children and adult FRP latencies and amplitudes across 5 key components of the FRP: P1 (neural), Nc (neural), n170 (neural), SP (artifact), and CR (artifact) (see Figure S3). For adults, not children, two sample t-tests indicated several key differences in FRP waveforms pre- and post-ICA; we observed significantly lower SP (*p* < 0.01) and CR (*p* < 0.01) amplitudes, suggesting that the ICA cleaning was highly effective at removing artifact. Further, we also observed significantly lower P1 (*p* < 0.01) and Nc (*p* < 0.01) amplitudes in adults, suggesting that the ICA cleaning also attenuated activity assumed to be of neural origin. Children showed no significant differences in FRP component amplitudes before vs after ICA cleaning (child P1, *p* = 0.7, child SP, *p* = 0.6, child Nc, *p* = 0.4, child n170 *p* = 0.5, child CR *p* = 0.7. Adult N170 was also not significantly different (*p* = 0.4). No frontal sources of activity remained after the removal of eye movement activity (Figure S2D), suggesting that global differences in theta if not driven by residual artifact are likely to originate in occipital visual processing areas (see also Figure 2 in the main text).

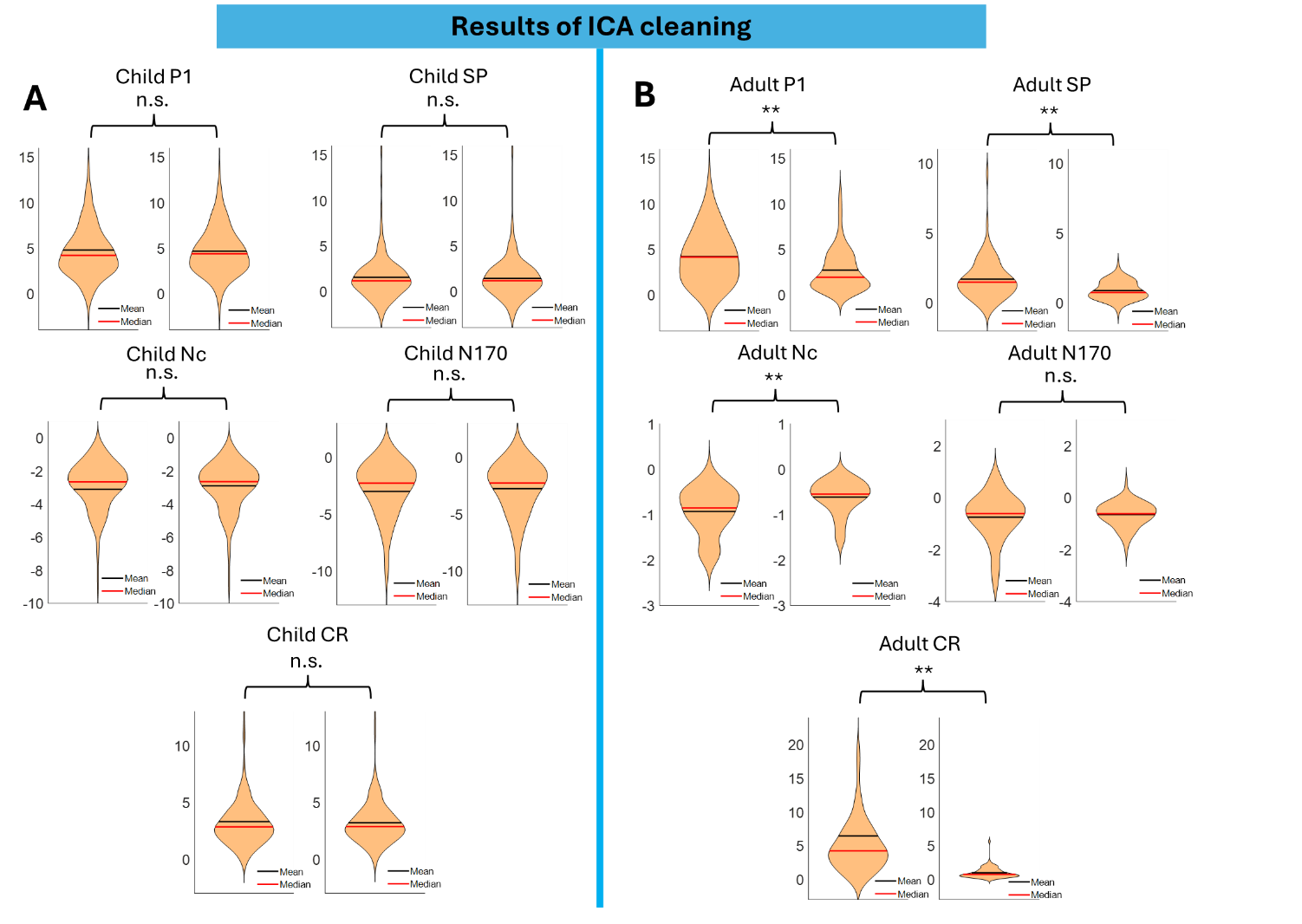

**Figure S3. Violin plots showing differences in peak amplitude of various FRP components in children and adults before and after correction using ICA.**

A. Shows comparisons of amplitude before and after ICA correction for 5 key components of child FRPs. P1 (occipital-neural), SP (frontal-artifact), Nc (central-neural), N170 (lateral occipital-neural), CR (frontal-artifact).
B. Shows comparisons of amplitude before and after ICA correction for 5 key components of adult FRPs. P1 (occipital-neural), SP (frontal-artifact), Nc (central-neural), N170 (lateral occipital-neural), CR (frontal-artifact).

** indicate the significance of p < 0.01 based on two-sample t-tests. ‘n.s’ refers to tests that were non-significant.

Overall, the ICA was more impactful on the adult data than the child data. Further, whilst these procedures did result in significant attenuation of the artifact, they also significantly reduced the amplitude of signals assumed to be of neural origin (see Figure 2 in the main text). Consistent with previous work, they were unable to attenuate eye artifact completely (Marriott Haresign et al., 2021). Overall, we felt that it was preferable to carry out the current analyses on data not cleaned using ICA.

3. **Time-frequency methods extended**

*Presentation of the bandpass filtered ERP*

Time-frequency representations of ERP amplitude and power often show broadband increases (Rousselet et al., 2007), that span the entire time range of the ERP waveform. Thus, in essence, the summed energy in the signal is measured across the various ERP components. This approach however can be insensitive to how different ERP components are differentially contributing to increases in spectral power. In this paper we use a less commonly presented output of time-frequency decomposition- the real result of convolution, essentially single trial ERP data that has been band passed filtered at increasing frequency bands from 1-40Hz. This is discussed in Mike Cohen’s book (Chapter 13, p 160). Unlike power, this value can be positive or negative depending on the phase relationship between the signal and the wavelet. In this paper, we have argued and shown evidence that visualisation and analysis of the data in this way allowed us to better understand the contributions of different components of ERPs/FRPs.

For example, in Figure 2 of the main text FRP activity related to eye movement artifact (labelled CR and SP) and neural processing of visual information (labelled P1) overlap heavily in time-frequency space. This is most evident in Figure 2C, images 1 and 3 of row C in which there are clearly observable increases in power over frontal and occipital electrodes which overlap significantly. When measuring power, this overlap makes it unclear what the source of the activity is. One possibility is this overlap is driven by propagation as the difference in peak latency between frontal and occipital signals is on the upper limit of what would be possible to observe due to the propagation of signals (Anderson, 2004). In other words, the increase in power could have been generated occipitally and propagated frontally. Another possibility is that these signals are the result of dipoles- with positive and negative activation across the scalp – i.e., that the increase in power could have been generated occipitally and been mirrored frontally. Another possibility is that they are independent signals generated by different neural systems/ processes. Disentangling these different potential explanations using power is highly speculative.

One more precise way to separate these signals would be to use ICA. However, these techniques are still quite limited in their ability to separate data into different sources even in very clean adult EEG data (Dimigen, 2020). Developmental data is accepted as being inherently noisier and even the most tailored approaches still cannot completely separate these sources of activity (Marriott Haresign, 2021). Thus, there is a need to consider other methods to separate these signals. The approach we are highlighting here gets around this issue primarily by representing positive and negative fluctuations in the waveforms. In Figure 2 (B), although both frontal pole and occipital activity very heavily overlap in time and frequency, it is clearer that these are two isolated sinusoidal signals and not the result of dipoles across the scalp. For example, in Figure 2B the positive and negative components of the frontal pole activity are matched in frequency and magnitude whereas the occipital activity has a subtle but noticeable difference in frequency and magnitude supporting the idea that these signals have independent sources. This is crucial as it allows us to be confident about what we are interpreting and analysing and neural activity (i.e., generated by the brain).

*Why power might not be the best measure (in this case)*

For developmental neuroscientists the approach described above offers several advantages over more popular approaches of examining spectral amplitude (absolute result of convolution) and power (amplitude squared). Firstly, it does not need to be normalised in order to visualise both low and high-frequency dynamics. One common way to do this is to use decibel conversion. The decibel (dB) is a ratio between the strength of one signal (frequency-specific band power) and the strength of another signal (a baseline level of power in that same frequency band). Where the ‘baseline’ refers to a period of time – typically a few hundred milliseconds before the start of the trial. This is not a problem for more traditional screen-based experiments which are designed such that there are sufficient breaks between stimulus onsets, to have a period of ‘rest’ to use as a baseline. However, when analysing free viewing data in children the stimulus onsets are driven by the participant's choice of when and where to fixate. The average fixation duration is 0.5s (with many, over 75th, lasting 0.2-0.8s), and each fixation onset produces signals in the EEG that last for several hundred milliseconds. This means that there is almost no point in the data in which the signal is totally at ‘rest’ which would serve as a suitable baseline for decibel conversion. We know the detrimental effects noisy baseline periods can have on decibel conversion. This is because signal amplitude was systematically underestimated in the noisier compared to the less noisy group (Gyurkovics et al., 2021).

4.
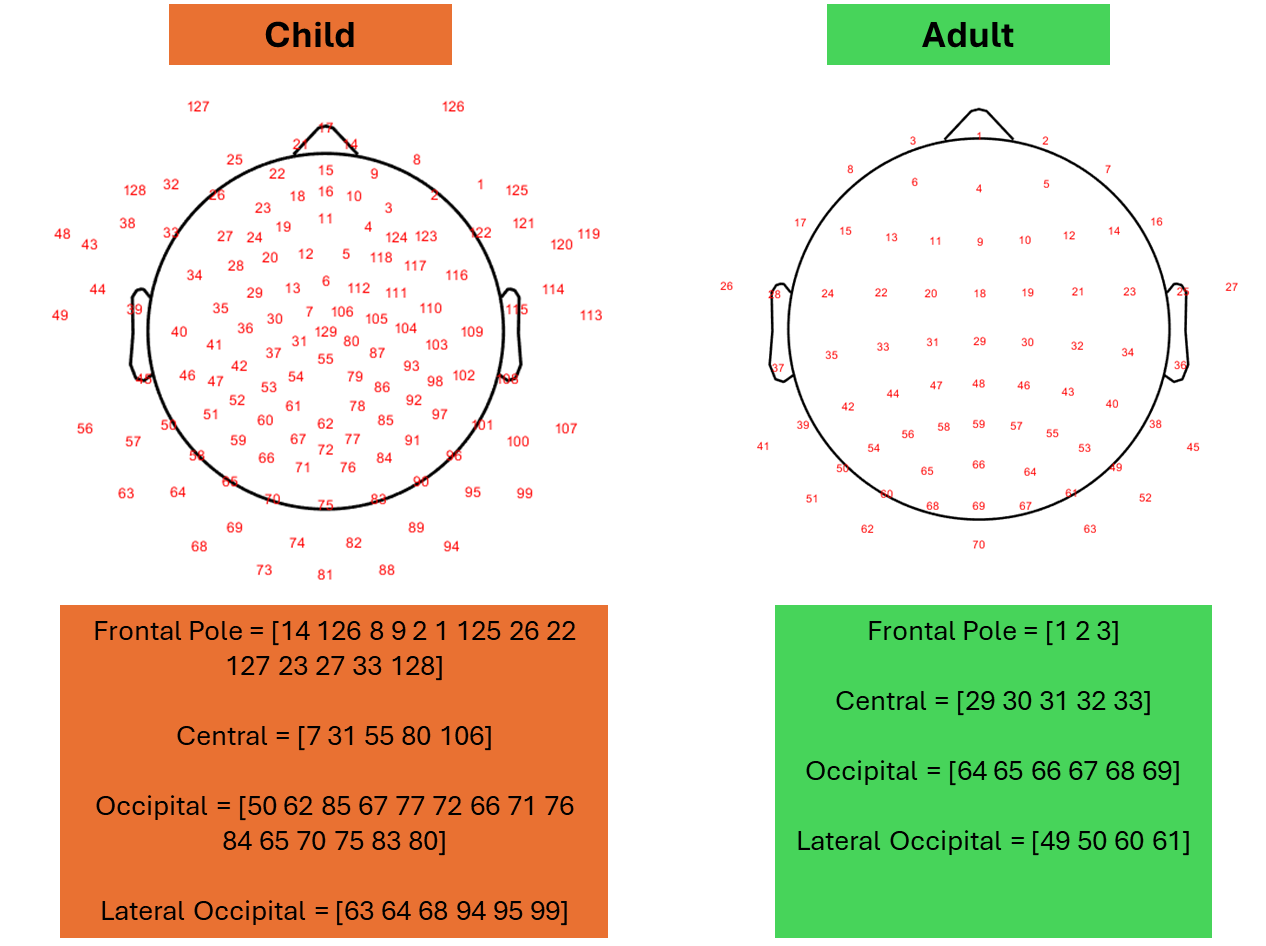
**Electrode groupings**

**Figure S4. Child and Adult electrode groupings**

5. **Adult theta differences across all data, FRP data and post FRP data**

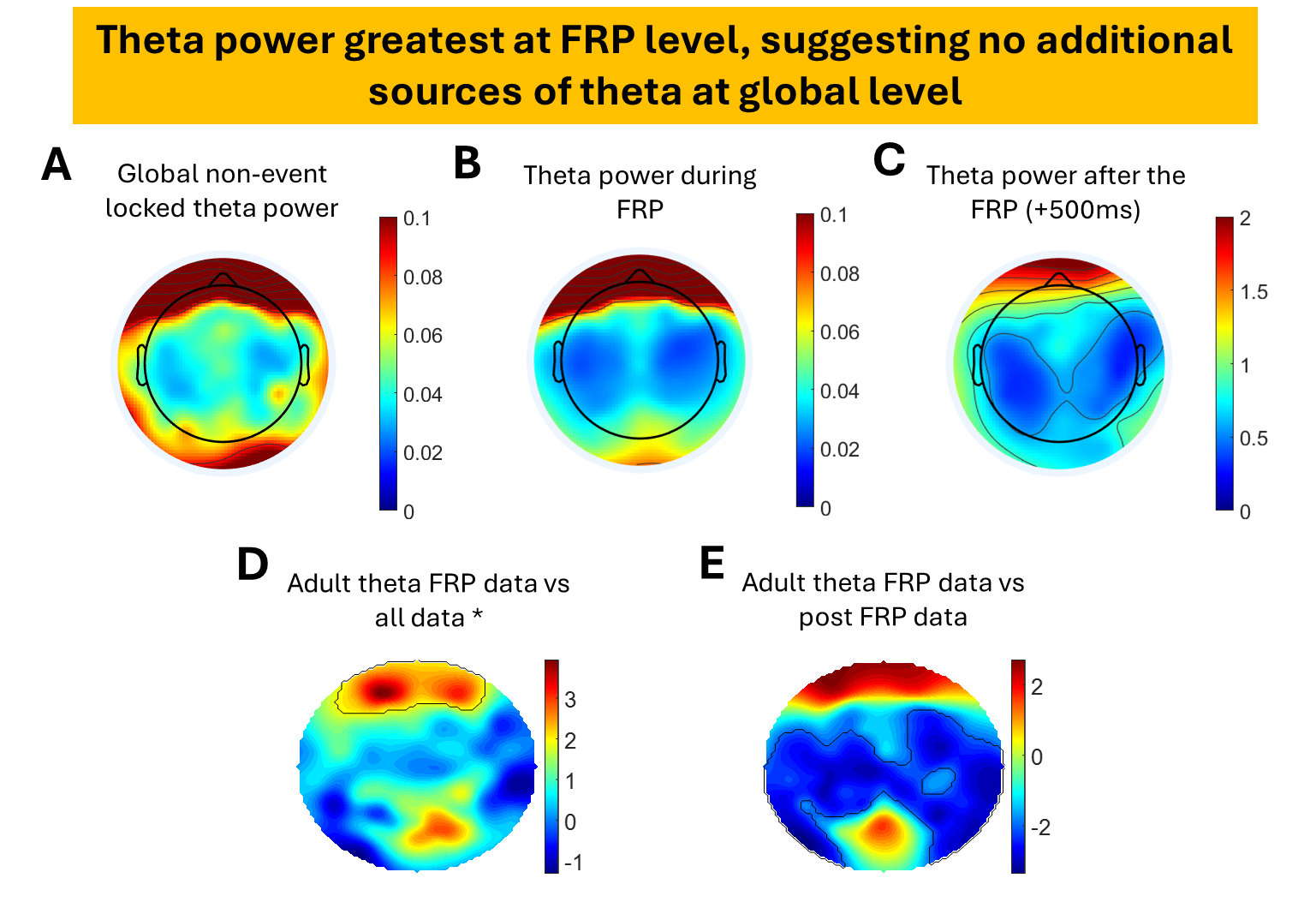

**Figure S5. Same as Figure 6, showing adult theta (4-8 Hz)** **differences across all data, FRP data and post FRP data. For all plots black outlines show significant clusters of increased power (p<0.05).**

A. Topographical distribution of adult theta power across entire data sample, not time locked to fixation onsets.
B. Topographical distribution of adult theta power during fixation related potential (FRP) time locked to onset of fixation.
C. Topographical distribution of adult theta power post FRP (+500 ms after fixation onset). A shorter post fixation window for adult’s vs children (+850ms) was selected due to shorter average fixation durations and Nc latencies.
D. Theta power during data segments where FRPs are and are not present.
E. Differences between adult theta for all data, FRP, and post FRP. Figure shows frontal and occipital theta is largely driven by activity of FRP as no sources of increased theta (in these areas) at global level (all data) and outside of the FRP.
6. **Inter-trial coherence (ITC)**

Figure S6 shows changes in amplitude, power, and inter-trial coherence (ITC) of child FRP activity over occipital and frontal pole electrodes. (The plots for amplitude and power are identical to those already shown in Figure 2 of the main text). Similar analyses exist for adult EEG data (Dimigen et al., 2011) and to provide a comparable picture of child FRP activity we plotted both occipital and frontal power and ITC (as did Dimigen). Overall, Figure S6 shows that increases in power associated with FRP activity (most notably P1 and SP) are closely associate with increases in ITC.  This reflects that, although power and phase estimations are thought to be independent, in reality, they are often difficult to separate (Burgess, 2013; van Diepen & Mazaheri, 2018). Changes in spectral power can give the appearance of increased phase locking/ resetting, due to changes in signal-to-noise ratios and errors associated with estimating phase (Muthukumaraswamy et al., 2011). This is relevant when discussing what combination of evoked and induced responses generates ERPs or FRPs (Burgess, 2012). The fact that we observe increases in spectral power and ITC makes it complicated to interpret what combination of evoked/ induced activity generated the FRP activity we observed in the present
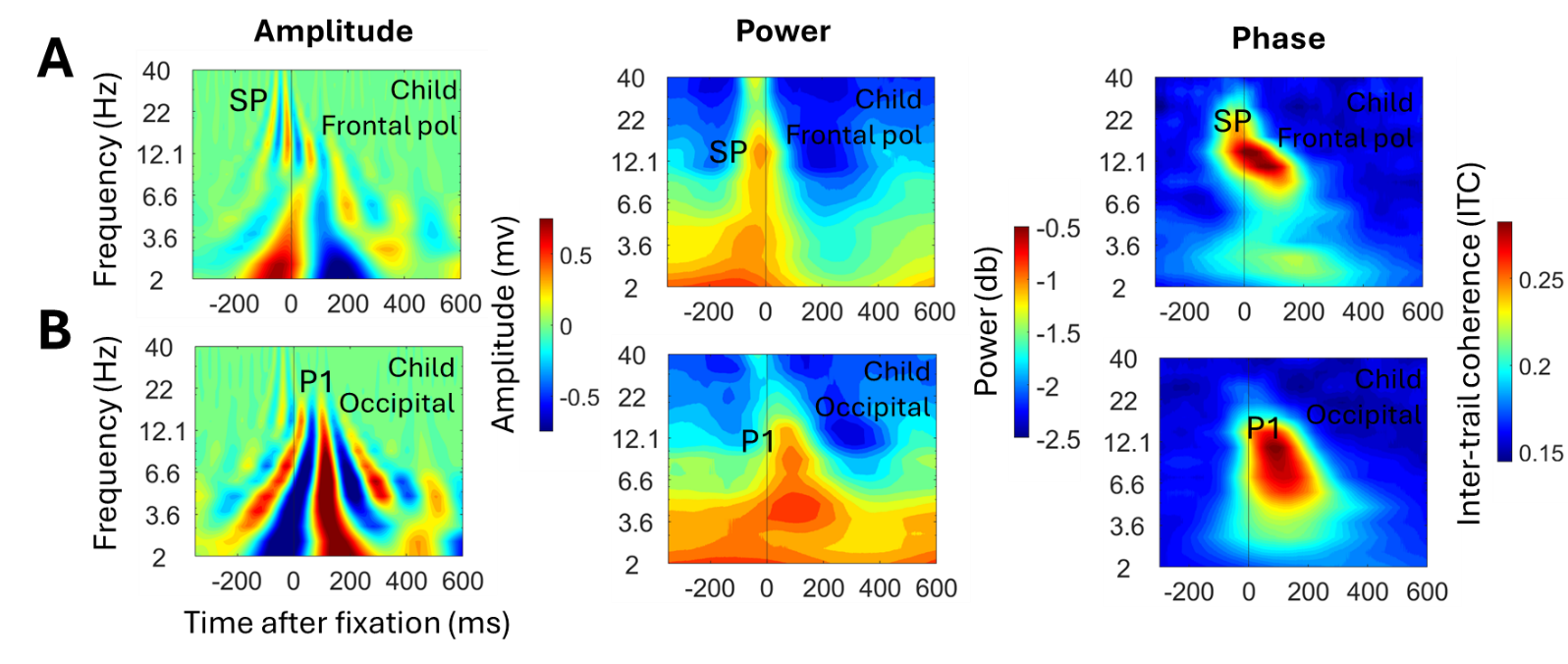
analyses (Sauseng et al., 2007).

**Figure S6. Amplitude, Power and Inter-Trial Coherence (ITC) of child FRP activity over occipital and frontal pole electrodes.**

A. Time-frequency Amplitude (same as Figure 2A in main text), Power (same as Figure 2C) and Phase (ITC) for child FRP activity over occipital electrodes

B. A. Time-frequency Amplitude (same as Figure 2B), Power (same as Figure 2C)

and Phase (ITC) for child FRP activity over frontal pole electrodes.

Additionally Figure S6A ITC plots support our findings in the main text (see section 3.2) that saccade-related activity manifests as lower frequency activity in children (above 10Hz, M = 14.1Hz) compared to adults (above 10Hz, M = 30.4).

7.
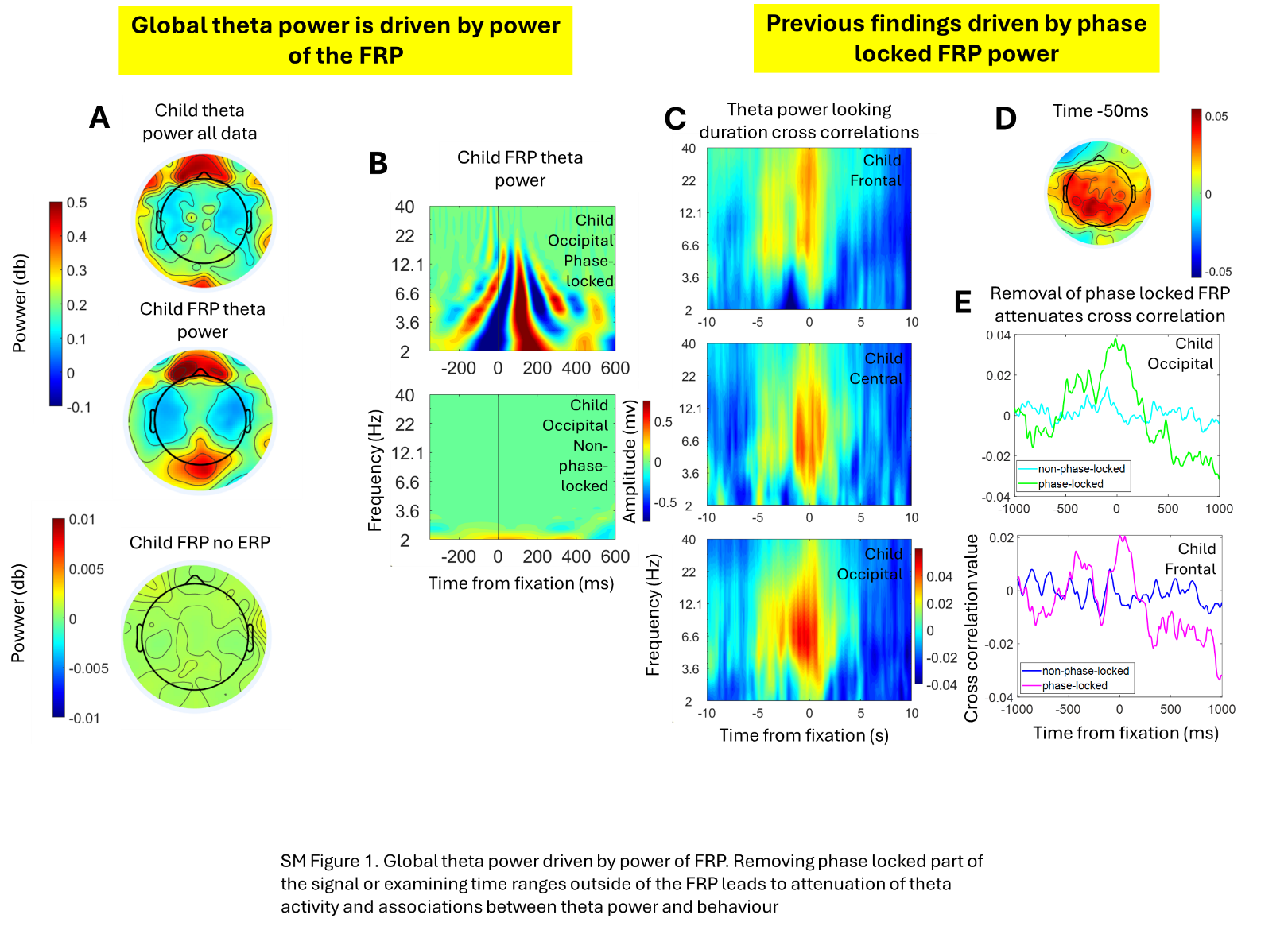
**Cross correlations between phase and non-phase locked power and fixation durations**

**Figure S7. Cross correlations between phase and non-phase locked power and fixation durations**

A. Topographical distributions of child theta power (2-7 Hz) across entire data sample, during FRP and after time range of FRP (+850ms)
B. Child time frequency amplitude plots of phase locked (with FRP) and non-phase-locked signals (FRP subtracted)
C. Child time-frequency cross correlations between power and fixation duration
D. Topographical distribution of cross correlation peak at -50ms
E. Child cross correlations between theta power (2-7Hz) and fixation duration before (phase-locked) and after (non-phase-locked) subtraction of the FRP.

We calculated child FRP theta power based both on the entire averaged data and on the averaged data after the subtraction of the non-phase-locked FRP (Fig S7A. Oscillatory activity was strongly attenuated after subtraction of the FRP, suggesting that changes in oscillatory activity are the result of additive changes (i.e. increased cortical excitability) time-locked to the offset of the eye movement (Cohen, 2014). Previous research (Perapoch Amado et al., 2023; Phillips et al., 2024; Wass et al., 2009) has observed time-locked associations between fronto-central theta power and infants’ attention to objects and people around them, which has been used to motivate claims that theta oscillations index active learning in infancy (Begus & Bonawitz, 2020). In Figure S6 we re-examine these associations. First (Fig S6C) we replicate these associations by using cross-correlations to show time-locked associations between theta power and attention episodes (in this case indexed by measuring fixations. Fig S6D shows the topographical distribution of the cross-correlation peak at -50ms. Then (Fig S6E) we repeat the analysis, concentrating just on the theta range (2-7Hz) after subtracting FRPs from the data. The previously observed time-locked associations are entirely removed. This suggests that the time-locked associations previously observed are attributable to fixation-related ERPs driving transient increases in theta power around the onset of a new attention episode.

8. **Inclusion/ Exclusion criteria**

*LEAP*. Exclusion criteria included substantial hearing or visual impairments not corrected by glasses or hearing aids, a history of alcohol and/or substance abuse or dependence in the past year and the presence of any MRI contraindications (for example. metal implants, braces, claustrophobia) or failure to give informed written consent to MRI scanning. The presence of co-occurring psychiatric conditions was not an exclusion criterion, given their prevalence in this population. Additionally, participants were excluded if they had a parent- or (where appropriate) self-report of a psychiatric disorder or had a *T*-score of 70 or higher on the self-report or parent-report form of the Social Responsiveness Scale-2. If on medication, all participants had to be stable (min. 8 weeks) at entrance point and over the course of the baseline visit to be included. Information on concurrent medication use was collected at the institute visit and substances were mapped to the Anatomical Therapeutic Chemical (ATC) classification system to categorise drugs as affecting/non-affecting the nervous system (ATC Level-1 code “N”; SM2.1).

*STAARS.* For full details see (Begum‐Ali et al., 2022). Participants were recruited for a longitudinal study running from 2013 to 2019. Infants could be enrolled in the study if they either had a first degree relative with ASD, a first degree relative with diagnosed or probable ADHD, or no first-degree relatives with either diagnosis. Information about diagnostic status was ascertained through a number of methods. Before families enrolled in the study, a telephone screening form was used to determine the presence of ASD and ADHD in family members. During their infant’s visit to the lab, the parent/caregiver also completed a “Medical and Psychiatric History Interview” with the researcher. The telephone screening form and this formal interview at a study visit were the primary sources of information about diagnostic status for either the parent or sibling. Typically, childhood diagnoses of ADHD were reported for older siblings, whilst parents were diagnosed with ADHD either in childhood or adulthood. In addition, we asked for medical updates at each study visit and re-administered the Medical and Psychiatric History Interview at the 2-year timepoint. We also requested diagnostic letters (pertaining to either the older sibling’s or the parent’s diagnosis) and asked parents to complete the DAWBA (Goodman et al., 2000) ASD and ADHD sections and these were reviewed by the senior clinician (TC). In addition, parents completed the Conners (Conners, 2008) (for ADHD) and the Social Communication Questionnaire (Rutter et al., 2003) and Social Responsiveness Scale (Constantino & Gruber, 2002) for ASD on the family member (either the older sibling or parent) with a diagnosis and where possible all other family members. This information is used to characterise our sample rather than for exclusionary purposes since, in the UK, NHS clinical diagnoses follow a gold-standard procedure including collation of information from parents, teachers and from in-person assessment that is beyond the scope of this study and more accurate than simple questionnaire measures.

A proportion of children/parents had suspected ADHD, but this had not yet been confirmed by clinical services. This is expected since we were targeting children with infant siblings, and often there can be significant delays in the diagnostic process for ADHD (e.g., (Auerbach et al., 2004, 2008). Further, up to 30% of children with ASD meet criteria for ADHD when prospectively assessed (Simonoff et al., 2008). In clinical practice, the prevalence of dual diagnosis is in practice much lower (Russell et al., 2014). Given the nature of the co-occurrence between ASD and ADHD and our longitudinal study, sometimes family members would have a suspected diagnosis of ADHD at study entry that would be confirmed later in the study; on other occasions, a family would enrol on the basis of an ASD diagnosis in an older sibling but by the end of the study, they would report that the same sibling was now undergoing assessment for suspected additional ADHD.

For those who reported suspected ADHD, screening questionnaires were used to examine the probable existence of ADHD. Inclusion decisions were reviewed by the project management team.

Specifically, for siblings (6 years or older) we used a shortened adapted version of the Conners 3 (Conners, 2008). Current behaviours that parents reported as occurring either “often” or “frequently” were scored. All included children met a minimum threshold for inclusion of i) 6 ADHD symptoms on either the hyperactivity/impulsivity scale (consisting of item numbers: 3, 43, 45[54]*, 61, 69[99]*, 71, 93, 98, 104) or the inattention scale (consisting of item numbers: 2, 28, 35, 47, 68[79]*, 84, 95, 97, 101), and ii) a positive score on the impairment scale (at least 2 out of 3 impairment items, consisting of item numbers: 106, 107, 108). Note that * indicates that these two items were collapsed into a single question in the adapted screening form.

For siblings aged less than 6 years, we used a shortened adapted version of the Conners Early Childhood (Conners & Goldstein, 2009). Behaviours that parents reported as occurring either “often” or “frequently” were scored. All included children met a minimum threshold for inclusion of i) 9 ADHD symptoms on the inattention/hyperactivity scale (consisting of item numbers: B8, B12, B22, B34, B42, B47, B49, B55, B65, B72, B74), and ii) a positive score on the impairment scale (at least 2 out of 3 impairment items, consisting of item numbers: IM1, IM2, IM3).

For parents, a shortened adapted version of the Conners Adults ADHD Rating Scale (CAARS; (Conners et al., 1999), either self or observer report. Current behaviours that parents reported as occurring either “often” or “frequently” were scored. All included parents met a minimum threshold for inclusion of 5 ADHD symptoms on either the hyperactivity/impulsivity scale (consisting of item numbers: 2, 4, 6, 8, 16, 18, 22, 25, 27) or the inattention scale (consisting of item numbers: 1, 9, 13, 14, 19, 21, 26, 29, 30). Of note, the adult version of the Conners does not include impairment questions.

Families who screened positive on this instrument were then included as a confirmed case However, it remains likely that within families with ASD, rates of actual ADHD are higher than those captured by our 1/0 diagnostically-based rating system.

9. **Multi-site data harmonisation**

Five sites acquired EEG data in LEAP: Kings College, London (KCL), The Central Institute of Mental Health, Mannheim (CIMH), University Medical Centre, Utrecht (UMCU), Radboud University Nijmegen Medical Centre (RUNMC) and University Campus Biomedico, Rome (UCBM). Three different EEG systems were used to acquire the data, Brainproducts Acticaps (KCL, CIMH, RUNMC), Biosemi Active-Two (UMCU) and Micromed (UCBM). Testing teams from each site attended initial training in London in 2013, followed by site visits to ensure correct set-up of equipment and that SOPs were followed. All sites then attended weekly telephone conferences to discuss data acquisition and quality, and to report any problems. Data were uploaded from each site to a central repository in their raw, manufacturer-specific, proprietary formats. Preprocessing and harmonisation of this data was performed at Birkbeck, University of London. Each dataset was first loaded into EEGLab. Briefly, the following steps were followed: 1) harmonisation of electrode labels to 62-channel common montage; 2) generation of horizontal electrooculogram (HEOG) channels from electrodes AF7/8 (KCL, RUNMC & UCBM only, CIMH & UMCU used external electrodes to record HEOG); 3) generation of variance-based data quality metrics and extraction of impedance values from Brainvision sites; 4) re-reference to FCz; 5) Resample to 1Khz; 6) harmonise event labels. This process resulted in harmonised data in a common EEGLab format, upon which all subsequent task-specific analyses were performed. Stimuli were presented using custom-written Matlab software (KCL, CIMH, UMCU, UCBM) and Presentation (UMCU).
